# Single-Cell Map of Diverse Immune Phenotypes in the Metastatic Brain Tumor Microenvironment of Non Small Cell Lung Cancer

**DOI:** 10.1101/2019.12.30.890517

**Authors:** Liang Wang, Jinxiang Dai, Run-Run Han, Lei Dong, Dayun Feng, Gang Zhu, Wei Guo, Yuan Wang, Min Chao, Ming-zhu Jin, Shi-Jia Jin, Dong-Ping Wei, Wei Sun, Weilin Jin

## Abstract

Cancer immunotherapies have shown sustained clinical success in treating primary non-small-cell lung cancer (NSCLC). However, patients with brain metastasis are excluded from the trials because the brain is viewed traditionally as an immune-privileged organ. The composition and properties of tumor-infiltrating myeloid cells in metastatic brain tumors are mostly unknown. To depict the baseline landscape of the composition, gene signature, and functional states of these immune cells, we performed - single-cell RNA sequencing (scRNAseq) for 12,196cellsafter data preprocessing, including 2,241 immunecells from three surgically removed brain lesions of treatment-naïve NSCLC patients. We found a lack of T lymphocyte infiltration and activation, as well as the vast expansion of tumor-associated macrophage(TAM) in the brain lesions of NSCLC patients. By comparing our scRNAseq dataset with published data from early and late-stage primary NSCLC tumors, we showed that this compromised T cell response is unique to brain lesions. We identified a unique alternative activation (M2) gene expression pattern of the TAM in the brain metastasis and a lack of known T cell co-stimulator expression. Accumulation of M2 polarized TAM may, therefore, cause the comprised anti-tumor T cell response in metastatic brain lesions. These findings can contribute to the design of new immunotherapy strategies for NSCLC patients with brain metastasis.

## Introduction

Metastatic brain tumors account for most lethality from central nervous system (CNS) malignancies in adults, which are approximately ten times frequent than a primary brain tumor^1^. The non-small cell lung cancer (NSCLC) attributes to almost half of the metastatic brain tumor diagnosed, followed by breast cancer, melanoma, renal cell carcinoma, and colorectal cancers ^2–4^. Patients treated with whole-brain radiotherapy (WBRT), surgical resection, and stereotactic radiosurgery often suffer disease relapse within five years after treatment ^5,6^. Although small molecule inhibitors targeting disease-associated driver mutations such as epidermal growth factor receptor(EGFR) or anaplastic lymphoma kinase(ALK) inhibitors in NSCLC have shown efficacy in controlling intracranial disease^7–9^, most treated patients eventual developed drug resistance^10–12^. Along with a large number of patients without actionable mutations makes new therapeutic options necessary^13,14^.

In the past five years, immune checkpoint therapies that target the T cell suppressive molecules PD-L1 and CTLA4 have significantly improved the outcomes in patients with advanced systemic disease for both NSCLC and melanoma^15–17^. In NSCLC, both nivolumab and pembrolizumab have been approved based on three significant randomized phases III studies^18–20^. A small number of patients with known active CNS disease enrolled in these trials; thus, no meaningful conclusions could be inferred regarding the impact of checkpoint blockade on the intracranial disease.

Several retrospective analyses of phase II have provided the preliminary support for a possible role of checkpoint blockade in treating active brain metastases^21,22^, and 3 clinical trials of ipilimumab in the treatment of metastatic melanoma reported an overall disease control rate (DCR) of 16% to 27% in patients with stable CNS disease^23,24^. In a recent retrospective analysis of five patients with new or progressing brain metastases from NSCLC treated with PD-1 blockade, an objective response was observed in two patients and persisted for greater than six months, suggesting a possible role for anti-PD-1 therapy in treating CNS metastases^25–27^.

While these findings are very encouraging, most PD-L1 expression positive primary NSCLC patients were failed to respond to conventional immune checkpoint therapy, let alone the ones without PD-L1 expression^28,29^.

The design of immunomodulatory strategies for the treatment of NSCLC patients with a metastatic brain tumor will tremendously benefit from a detailed understanding of the immune cell landscape that develops specifically in response to tumor cues^30^.

To shed light on the complexity of tumor-infiltrating immune cells in NSCLC with a metastatic brain tumor, we performed deep single-cell RNA sequencing on 12,196 cells, including 2,012(20.4%, 95% CI: 4.1% - 36.7%) myeloid cells from 3 surgically removed brain lesions of treatment-naïve Lung adenocarcinoma (LUAD)patients. We found that lack of T cell infiltration and activation in the brain metastasis of LUAD as well as triple negative breast cancer brain metastases and glioblastoma. We identified a unique signature of the tumor-associated macrophage (TAM) in the brain metastases compared with the early and late-stage primary LUAD. We also did bulk RNA analysis on brain metastases from additional 6 LUAD patients to supplement to the limitation on low number on patient sample with single RNAseq. The low inflammatory nature of tumor cells in brain lesion and lack of T cell co-stimulator expressed on the antigen-presenting cells is one of reason underneath the lack of T cell activation.

## Results

### Immune landscape of brain metastasis of lung adenocarcinoma

To generate a transcriptional map of the immune cell states

In human brain metastasis of lung adenocarcinoma, we constructed an atlas comprising 12,346 cells collected from freshly resected tumors of 3 treatment naïve patients (supplemental table 1). A total of 23,467 cells were subjected to single-cell RNA sequencing (scRNAseq) using the 10X genomics inDrop platform, with an average depth of 7,778 unique molecular identifiers (UMI) counts per cell(Supplemental fig.1).

Using tSNE to visualize high-dimensional data in two dimensions while preserving single-cell resolution^31^, we analyzed the distribution of the different stromal cell lineages that accumulated in metastatic lesions. These identified cell clusters that, through marker genes, could be readily assigned to known cell lineages: in addition to cancer cells, we identified immune cells (myeloid, T, and B cells), oligodendrocytes, endothelial cells, and pericytes (Figure 1A,B).

**Figure1.**
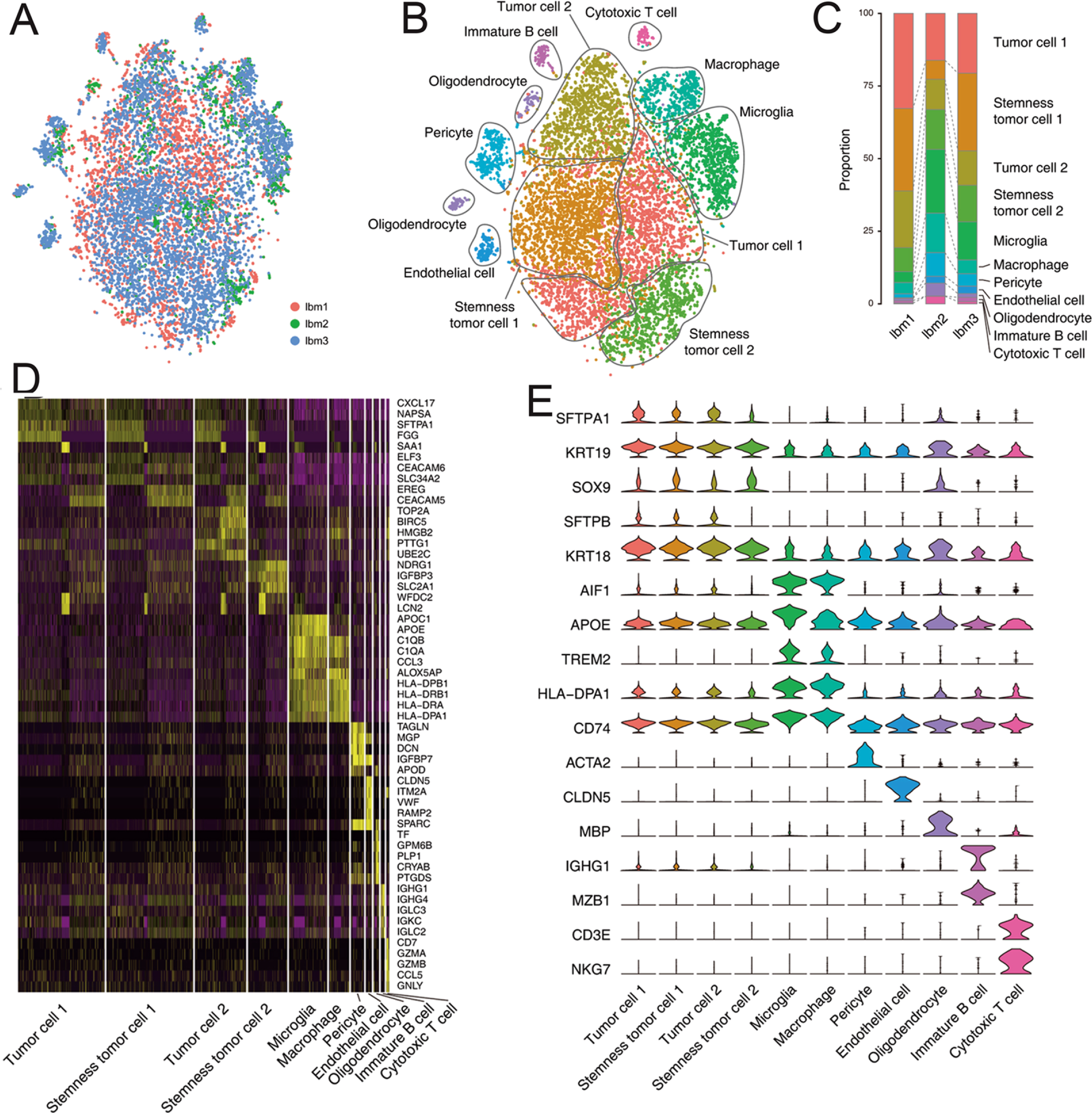
Immune landscape of brain metastasis of lungadenocarcinoma. (A, B) t-Distributed stochastic neighbor embedding (tSNE) of the 12,196 cells profiled here, with each cell color-coded for the corresponding patient(A) and the associated cell type (B). (C) Proportions of major celltypesfor each lung adenocarcinoma (LUAD) patient’s tumorinfiltrating immune cells, colored by cell type. (D) Heatmap shows scaled average expression levels of top5 differentially expressed genes (DEGs) for eachcell-type.Top5 DEGs for each cell type are determined byhigher expressional fold-changes against all others. Highest expression is shown inyellow and lowest in purple. Select lineage-specific genes are shown on the right. (E) Violin plots show the expression of marker genes for each cell type.

We first verified that major immune cell types were identifiable in each patient using canonical markers (Methods). The tumor-resident immune cell compartment comprised mostly of mononuclear phagocytes, including microglia (AIF, APOE, SPP1, TREM2, CSF1R, C1QA) (Gosselin et al. Science.2017) and monocyte derived macrophage (CD14, CD74,CD68, HLA-DR) (Figure 1D,E). In contrast, T and B lymphocytes only account for10.3%(95% CI: 6.9% - 13.7%) of the total immune cell population. Strikingly, the other major myeloid population: dendritic cells, neutrophils, and NK cells, are absent in the metastatic brain lesion (Figures 1B).

To validate our single RNAseq with brain lesions of LUAD patients, we did bulk RNA sequence on frozen human autoptic tissue specimens of brain metastases from 6 more LUAD and analysis their immune composition using CIBERSORT software^32^. Consistent with our finding with scRNAseq, the absence or low presentation of neutrophil and expansion of mononuclear phagocytes in brain lesion is confirmed (supplemental fig.2A).However, there are more predicted T/B/NK cell population than we found in scRNAseq samples, which could reflect the tumor microenvironment variability across patients or lack the accuracy of CIBERSORT in predicting certain types of immune cells in complex tissue^33^.

#### Conclusion

We found a lack of T and B lymphocyte infiltration, as well as the vast expansion of tumor-associated macrophage(TAM) in the brain lesions of LUAD patients.

### Immune microenvironment of the brain metastasis is unique compare to primary tumor

Lack of T lymphocyte infiltration and vast expansion of tumor-associated macrophage (TAM) in the brain lesions of LUAD patients is a striking contrast to related abundant of lymphocyte infiltration in primary NSCLC patients as shown in previous studies^34^. We validated these findings by histology analysis of T cell (CD3+), B cell(CD20+), and macrophage(CD68+) in both primary and brain metastatic lesions of LUAD patients (supplemental fig.3). We analyzed 19 human tissue specimens of brain lesions from Lung adenocarcinomapatientsand a tissue array contains 52 primary NSCLS samples (supplemental table1). Profound microglia/macrophage activation with marked peritumoral accumulation and some intra-tumoral infiltration of CD68-positive microglia/macrophages was found as previously described^35,36^. Only a few B- and T-lymphocytes were observed in and around BM,with tertiary lymphoidstructures (TLS) (supplement fig.3), consistent with our findings by single cell RNAseq.

The stromal cells in tumor microenvironment has known to regulate the T cell response in various tumor^37^, we first ask whether lack of T cell infiltration in brain metastases in LUAD patients is due to its unique stromal cell status.To gain insight into the molecular signature of the tumor stromal cells across multiple stages of LUAD progression, were-analyzed published scRNA seq data sets from early-stage LUAD ^38^and late-stage LUAD^39^, along with our brain metastatic lesions (Method). Single-cell profiles of non-malignant cells highlighted the composition of the tumor microenvironment. We normalized and analyzed the distribution of non-malignant cell lineages that accumulated in early-stage, late-stage primary, and brain metastatic LUAD across patients. We annotated clusters by the expression of known marker genes as T cells, B cells, plasma cells, NK cells, macrophages, dendritic cells, mast cells, endothelial cells, pericytes, and oligodendrocyte, astrocytes. Consistent with previous findings, we also found T and B lymphocytes were present at a higher frequency in the tumor microenvironment in both early (55.8%), and late-stage LUAD (83.2%) compared to brain metastasis (10.3%)(Figure 2A,D).In contrast, the proportion of mononuclear phagocytes was not changed much between primary lung tumor and metastatic brain lesions of LUAD patients(Figure 2A,D). The marked transcriptional change was observed in majority of the stromal cell types based on their stage and tumor sites, visualized by distinct separation of plots in tSNE (Figure 2B,C). The stromal cells from metastatic brain lesions of LUAD are unique by their gene expression profiles, visualized by distinct spatial separation of early-stage, late-stage, and brain metastatic stromal cells in the principal component analysis (Figure 2C).

**Figure2.**
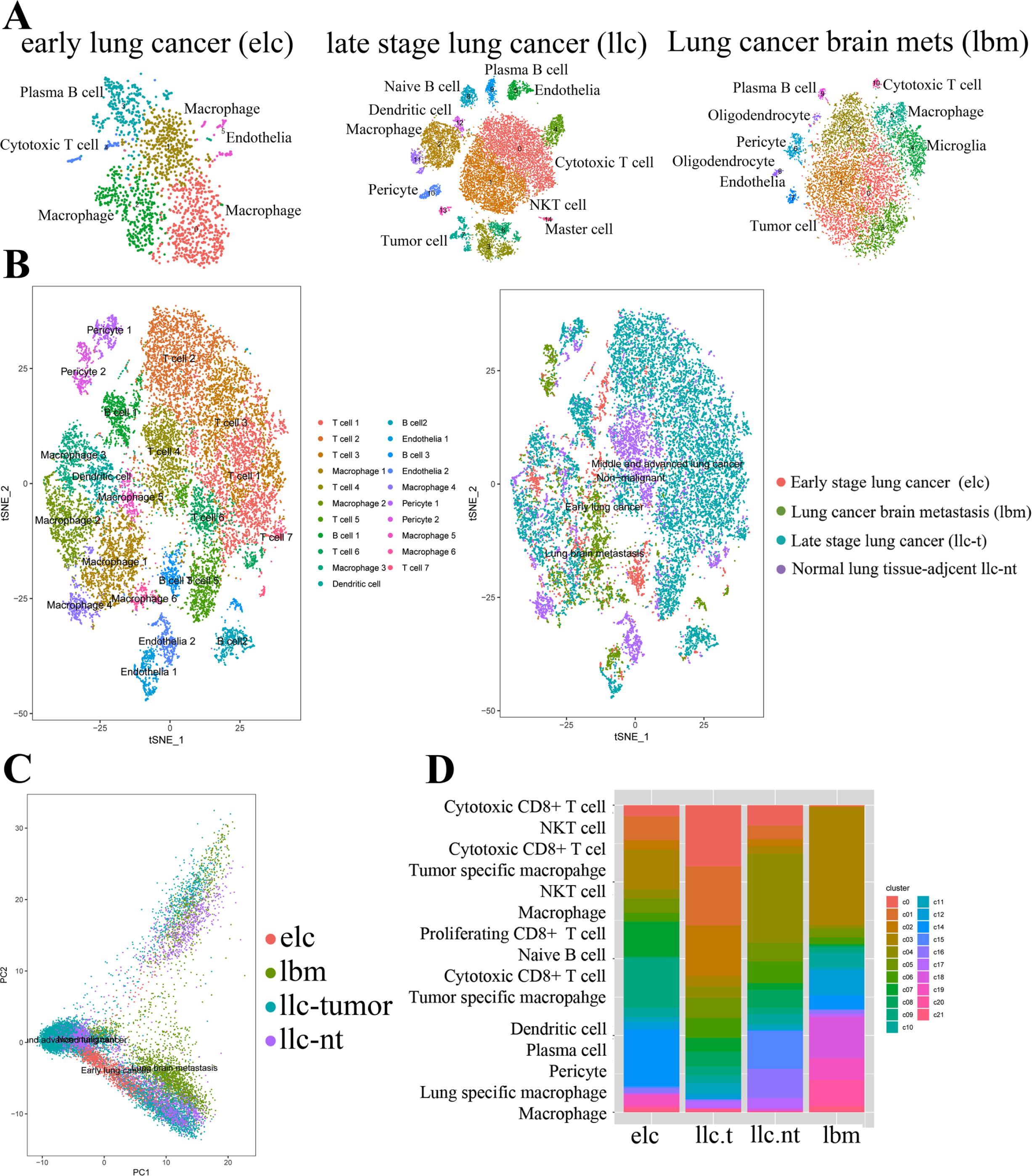
Immune microenvironment of the brain metastasis is unique. (A) tSNE of 1,920 cells from early stage LUAD (elc) (Lavin,et al. Cell) and 18,800 cells from later stage LUAD (llc)(Lambrecht, et al. Nature Med), along with 3,082 cellsfrom brain lesions of LUAD (lbm). (B) tSNE plot of combined 20,089 stromal cells from LUADs at different stages (as A), color-coded by their associated clusters (left) or the sample type of origin (right).(C) Principle component analysis (PCA) based on transcriptomes of single stromal cells from different stages of LUAD patients.(D) Proportions of major celltypes for each cluster derived from (B) as combined all tumor-infiltrating immune cells from all three different stages of LUADs, colored by cell type.

Myeloid cells have a unique ability to control T cell function at the tumor site, through their ability to present tumor-associated antigens to T cells and to produce critical T celldifferentiation cytokines^40–42^.Since the number of macrophage did not change across LUAD at different stages, we asked whether the molecular composition in primary tumor could be different from that in metastatic brain tumor.First, by mean of gene expression, the tumor specific macrophage clusters in primary tumor and brain lesion were not different a lot (Figure 2B). We found that beyond mean expression levels, covariance parameters varied significantly between these macrophage cluster (Figure 3A). This occurs due to the genes typically being co-expressed in the same cells in one cluster (positive covariance) but expressed in a mutually exclusive manner in the other cluster (negative covariance)^43^.

**Figure 3.**
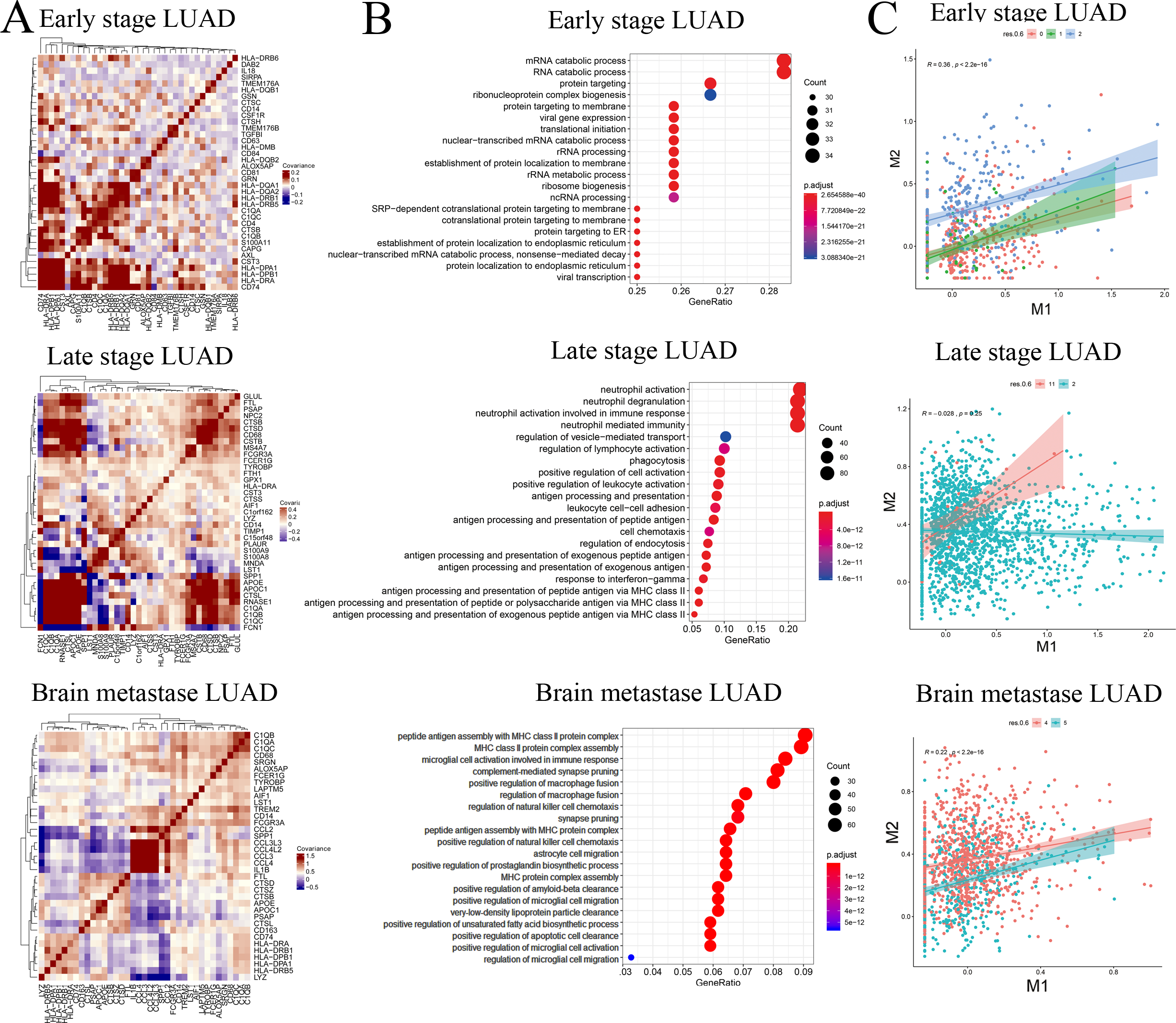
Unique macrophage profile in the brain metastasis reveals its alternative activation of M2 signatures. (A) Scatterplot of normalized mean expression of M1 and M2 signatures per cell (dot); cells assigned to TAM clusters have been highlighted by colors. (B) Heatmaps showing covariance efficiencyofexpressionof selected macrophage marker genes in three TAM clusters. (C) Gene ontology (biological process, BP) enrichment analysis of DEGs in TAM in different stages of LUADs (elc, cluster 1; llc, cluster 2; lbm, cluster4). The graphs show top20over-represented GO groups withhigher GeneRatio of DEGs. The size of each bubble reflects the number (Count) of DEGs assigned to eachGO BP term. Color of the bubbles displays adjusted p-value.

We also found distinct pathway expression levels of tumor-associated macrophage in brain metastasis vs. early and late-stage LUAD (Figure 3B), we noticed a strong enrichment of peptide antigen assembly with MHC protein complex in the metastatic brain lesion (Figure 3B bottom panel). We noticed a substantial increase of inflammatory with neutrophil mediated immunity in late stage LUAD (Figure 3B middle panel). In early-stage LUAD, the mRNA catabolic process and ribonucleoprotein complex biogenesis are enriched, indicating an anti-viral like immune response in early stage lung tumor(Figure 3B top panel).

Furthermore, expression of genes associated with ‘‘alternatively activated’’ (M2) macrophages, including scavenger receptor CD163, Mannose Receptor MRC1, and inhibitory molecule B7-H3 (CD276)^44^, increased in brain metastatic lesion (Figure 3C). Concomitantly, immunostimulatory genes associated with ‘‘classically activated’’ (M1) macrophages, including chemokine CCL3/CCL4 (MIP-1a), also increased in brain metastatic lesion (Figure 3A, supplemental fig.2B). Among all TAM populations, particularly TAM clusters in brain lesion (cluster 4), were among the monocytic clusters with the highest expression of the canonical M2 signature but were likewise high in the M1 signature (Figure 3C). Quite strikingly, we found that M1 and M2 gene signatures positively correlated in the TAM populations in early stage LUAD (cluster1) and brain metastatic lesion (R=0.36 and R=0.26 respectively), but TAM in late stage lung tumor (cluster 2) more towards to M1 polarization instead (R=0.03) (Figure 3C), in line with recent findings in other tumor types^45^.

#### Conclusion

The stromal cells,largelymacrophage,in brain metastases are molecularly distinct from their counterpart in both early and late stage primary LUAD.

### Paired immune cell mapping reveals a distinct macrophage signature in brain metastases of Lung Adenocarcinoma

To fully capture the heterogeneity of the TAM compartment, we first performed an unbiased single-cell transcriptomic analysis of myeloid cells that accumulated in early-stage, late-stage primary tumors and brain lesions (method). This analysis revealed seven myeloid cell clusters distinguished based on characteristic gene expression profiles identified according to highly expressed and differential genes (Figure 4A). Specifically, we identified a dendritic cell (DC) cluster expressing high levels of MHC class II molecules, CD1C, FCER1A, CLEC4A, and CD207 (Figure 3B, clusters 10), which is found in primary lung tumor but absent in metastatic brain lesion. Other clusters(3, 5, 9,16,19,20) were identified as macrophages (Figure 4A,B) based on the differential expression of CD68, CD163, and CSF1R (Figure 4B). Despite this diverseness, tSNE plots showed rather poor separation of clusters, suggesting that they represent diverse cell states on a graded scale rather than separate entities, in line with the spectrum model of macrophage activity^46^. Strikingly, tSNE plotting revealed a dichotomy between early-stage, late-stage primary tumor or brain lesion derived macrophages (Figure 4C).

**Figure4.**
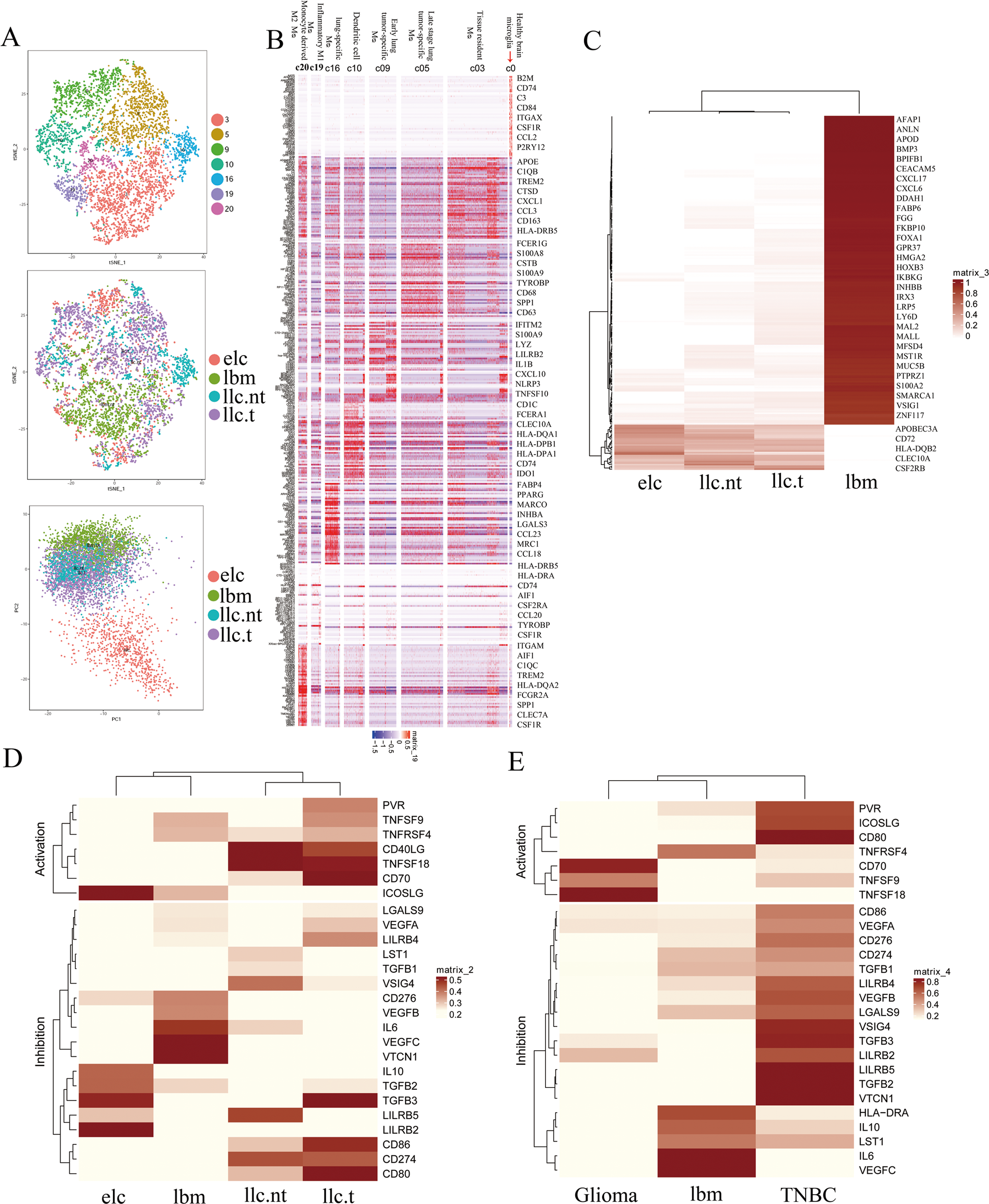
Molecular profile of the macrophage in different stages of lung adenocarcinoma. (A) tSNE and PCA plots showing5,187 myeloid cells from different stages of LUADs (elc, llc vs lbm). Color-coded by their associated clusters (top) or the sample type of origin (middle and bottom). (B) Heatmap showing DEGs in each cluster of TAMs cellsfromdifferent stages of LUAD(elc, llc, lbm). Columns represent single cells with lowest expressed genes in blueandhighestinred. Select phagocytes-specific genes are shown on the right. (C) Differential gene expression analysis reveals uniqueTAM signature in brain metastasis of LUAD. (D) Heatmap showing expression of distinct T cell co-stimulatory and co-inhibitory ligands on macrophages from brain metastases of LUAD (lbm) versus primary lung cancers (elc, llc) (left), and also from brain lesions from LUAD (lbm), breast cancer brain metastases (TNBC) and glioma (right).

With 2,012myeloid cells detected after data processing, microglia/macrophage represents the most prevalent stromal cell type in metastatic brain lesions. The tissue-resident macrophage population was identified as distinct cluster 3, which composed macrophages from primary lung tumor and metastatic brain lesions, with distinct gene expression AIF1, APOE, TREM2, SPP1, CD68, CD63(Figure 4B). Enrichment of AIF1, APOE, TREM2 has been shown as the unique feature of microglia^47^, the tissue-resident macrophage in the brain, which is also highly expressed in residual macrophage,e.g.,Langerhans cells in the lung^39^.One macrophage cluster comprised predominantly of cells from metastatic brain lesions (cluster 20) were distinct from those comprised predominantly of cells from early-stage tumor lesion (cluster 9) and late-stage tumor lesion (clusters 5). This resultindicates that tumor-associated macrophages in brain lesions were distinct from their primary tumor and tissue-resident macrophage counterparts.

The early stage lung tumor-associated macrophages up-regulated the pro-inflammatory S100 calcium-binding protein A8/A9 (S100A8/A9), IFITM2, IL1B, and NFKBIA (Figure 4B). A similar pro-inflammatory profile was found in TAM in late-stage lung tumor lesion, with additional up-regulation of CCL2 and VEGFA. In contrast to the primary tumor, the macrophage in brain lesions shows a more antigen-presenting profile with the up-regulation of various MHC II molecules (HLA-DRA, HLA-DQA, HLA-DPB) and also a component of the complement (C1QA, C1QB). This likely reflects differentiation and activation of either recruited or tissue-resident macrophages.

When different pathway expression levels of tumor-associated macrophage in brain metastasis vs. early and late-stage LUAD (supplemental fig.4), we noticed a strong enrichment of peptide antigen assembly with MHC protein complex across the evolution of the tumor. In early-stage LUAD, the positive regulation of regulatory T cell differentiation and Fc receptor-mediated inhibitory signaling pathway are enriched(supplemental fig.4A), indicating T regulatory cell may play a role in tumor immune-editing at an early stage consistent with the finding previously^38^.In the late stage, LUAD, neutrophil aggregation, and production of immune inhibitory cytokines are more prominent, such as IL10 and transforming growth factor beta1 production(supplemental fig.4B). In contrast, in the metastatic brain lesion, complement-mediated synapse pruning, positive regulation of microglial cell activation, and neutrophil clearance, indicating a brain-specific anti-inflammatory response to the tumor(supplemental fig.4C).

#### Conclusion

Detailed analysis of the macrophage activation in the brain lesion of LUAD patient versus its primary tumor counterpart reveals its unique polarization states. The antigen presentation process is not disrupted in macrophage in brain lesion.

### Molecular profile of the macrophage in a different stage of lung adenocarcinoma reveals brain-specific TAM lack of conventional T cell co-stimulatory

Since the antigen presentation process is not disrupted in brain metastases of LUAD patients, we went on to test whether tumor cell intrinsic factors are part of the cause of lack of T cell in brain lesions. We analyzed bulk RNA sequencing data from 14 early stage and 11 late stage primary LUAD tumor from TCGA database (supplemental table 1), and alsobrain metastasis lesion from additional six treatment naïve patients. An unsupervised cluster of genes that overexpressed in tumor cells of metastatic brain lesions are distinct from the primary tumor(supplemental fig.5A). However, PCA analysis showed the early stage (elc) and late-stage LUAD (llc) tumor cell gene expression is more closeto each other than brain metastases(supplemental fig.5B). When contrasting pathway expression levels in brain metastasis vs. early and late-stage LUAD, we noticed a substantial reduction of I−kappaB kinase/NF−kappaB signaling, interleukin−1−mediated signaling pathway and antigen processing and presentation of peptide antigen via MHC class I in brain metastases (supplemental fig.5C). In contrast, purine ribonucleotide synthesis, oxidative phosphorylationand regulation of metallopeptidase activity are up-regulated in metastatic brain lesions(supplemental fig.5D). These data support low inflamed features of tumor cells in metastatic brain lesions but not in primary tumors.

The fact that tumor-associated macrophage composed most of the myeloid-derived suppressive cell population, it is much likely that lack of lymphocyte infiltration in metastatic brain lesion may attribute to the unique immune-suppressive feature or lack of immune co-stimulatory factors in myeloid cells in brain lesions in comparing to primary tumors.

To fully capture the immune-modulatory feature of the TAM compartment, we performed an unsupervised cluster of different gene expression of known immune-stimulatory vs. inhibitory molecules in myeloid cells that accumulated inearly-stage, late-stage primary tumor and brain lesion of LUAD patient (supplemental table 2). The expression of ligands for immune checkpoint molecules, including approved targets CD274(PD-L1) and CD80 (ligand of CTLA4), but also others currently targeted in clinical trials ICOSLG (ICOS/CD278), TNFRSF9(4-1BB/CD137), CD70/TNFSF7(CD27), HLD-DRAs

(LAG3), TIGIT (CD155), LGALS9 (HAVCR2/TIM3) among others are drastically different between lung tumors at different stages(Figure4D). Both T cell stimulatory and inhibitory factors are highly expressed in late-stage lung tumors, which is consistent with the abundance of cytotoxic and exhausted T cells in this tumor^48^. However, macrophage in both early-stage lung cancer and brain lesions are lack of ligands for T cell stimulatory and conventional inhibitory factors (CD274, CD86)(Figure4D). This is consistent with the lack of exhausted T cells in both early-stage lung tumors and metastatic brain lesions.

Instead of enrichment of T regulatory cells in early lung tumors^38^, the brain metastatic lesion overexpression unique set of T cell inhibitory factors including CD276/B7-H3, VTCN1 (ligand for CD272/BTLA), VEGF, IL6, which is expressed in primary lung tumors. CB7-H3, VTCN1, VEGF are also highly expressed in macrophage from breast cancer brain metastatic lesion but interestingly is absent from advanced glioblastoma(Figure4D,E).

#### Conclusion

These results confirmed the non-inflamed (cold) feature of brain lesion of lung adenocarcinoma. The macrophage in their microenvironment also lack expressionofT cell co-stimulatory factors.

### Unique immune cell profile in brain metastasis comparing with glioblastoma

We found both metastatic brain lesions from lung and breast cancer brain metastasis sharing the expression of T cell inhibitory factors but not the glioblastoma. Comparing the cellular and molecular profiling of metastatic brain lesion (lung and Breast) with the primary tumor of brain (glioblastoma) provide a great opportunity to understand to what extent the tumor microenvironment is shaped by the tumor of the origin or the location of the tumor.

To compare the molecular signature of tumor stromal cells from brain metastases of lung adenocarcinoma and breast cancer with that of glioblastoma, we analyzed6,000 cells from freshly resected tumors from brain lesions of treatment naïve triple-negative breast cancer patient. We re-analyzed published scRNA seq data sets from 38 advanced glioblastoma patients^49^. We normalized and analyzed the distribution of non-malignant cell lineages that accumulated in these three types of brain tumors. We annotated clusters by the expression of known marker genes as T cells, plasma cells, NK cells, macrophages, dendritic cells, mast cells, endothelial cells, pericytes,oligodendrocyte and astrocytes(Figure 5A).

**Figure 5.**
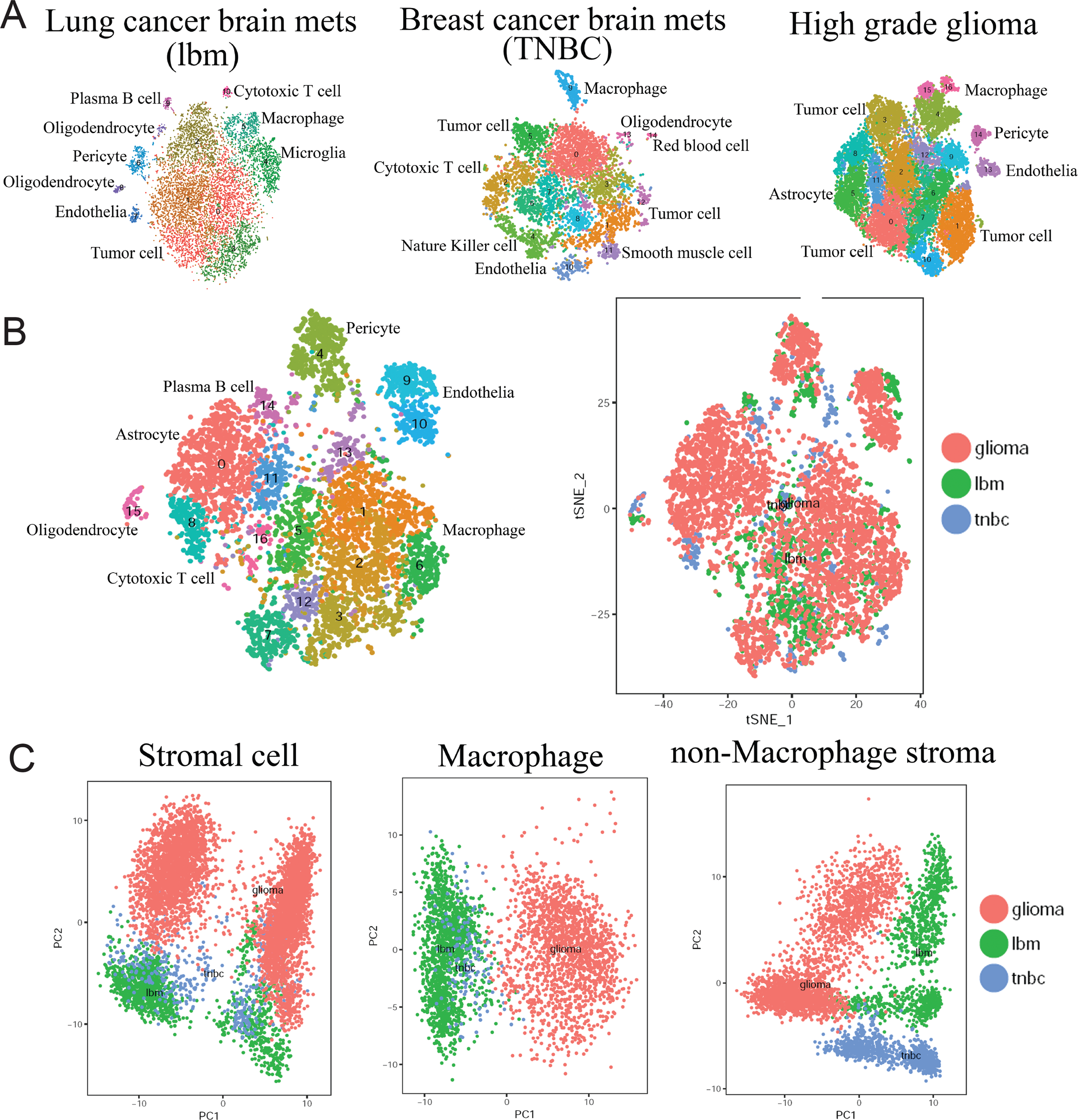
Unique immune cell profile in brain metastases comparing with glioma. (A) tSNEshowing clusters of single cells from resected lesions of LUAD brain metastases (3X patient, 12,196cells), triplenegative breast cancer brain metastases (1X patient, 6,000 cells) and highgrade glioma (38X patients, Yuan J, et al. 21,994cells). (B) tSNE showing clusters of stromal cells from three types of brain tumors(lbm, TNBC, glioma). (C)PCA analysis of stromal cells, macrophages and no-macrophage cells from the brain metastases of LUAD,TNBC and high grade glioma.

Consistent with previous findings and in line with our findings in LUAD brain metastasis, we also found T and B lymphocytes were almost absent in the breast cancer brain Mets and utterly absent in glioblastoma. The mononuclear phagocytes wereabundant and account for majority of stromal cells in all three types of brain tumors. The marked transcriptional difference was observed in the majority of the stromal cell types based on their tumor of origin, visualized by distinct separation of plots in tSNE (Figure 5B,C). The stromal cells from metastatic brain lesions of LUAD and breast cancer are clustered together, separate from the counterpart from glioblastoma by principal component analysis (Figure 5B,C). Macrophages from metastatic brain lesions of LUAD and breast cancer are transcriptionally moreclose to each other than glioblastoma despite their different tumor of origin. These data indicatetumor cells that metastasize to the brain from peripheral organs elicit a different but similar innate immune response compared with tumor origin from the brain, e.g.,glioblastoma. These data emphasize the impact of tumor of origin in shaping the tumor immune response rather than the local microenvironment.Strikingly, the non-macrophage compartment of the stromal cells (endothelia, pericyte, astrocyte, oligodendrocyte) are mostly separate from each other of these three tumor types, further supporting the dominant role of cancer cell-intrinsic mechanisms in shaping the tumor microenvironment^50^.

#### Conclusion

When comparing with the tumor arising from brain, e.g., glioblastoma, molecular signature of stromal cells derive from lung and breast brain metastases are very distinct from their counterpart in glioblastoma. The tumor immune microenvironment of brain metastases is shaped by tumor intrinsic property that being inherent of their organ of origin (Lung and breast).

### Clonal expansion of B and T cells in brain metastases of LUAD patients

T and B cell receptor (TCR and BCR, respectively) Vβ or immunoglobulin heavy chain complementarity-determining region three sequencing allows monitoring of repertoire changes through recognition, clonal expansion, affinity maturation, and T or B cell activation in response to antigen^51^. Although the number of B and T cell are relatively lower in metastatic brain lesion of LUAD, the B cell appear to be mature plasma that high express immunoglobulin G and T cells are mostly composed of cytotoxic T cells (Figure 1D,E). Importantly, many patients include the three patients we had performed scRNA seq, had tertiary lymphoidstructures (TLS), which accumulated near the invasive tumormargin or within the tumor mass that was absent from healthy lung (supplemental fig.2C).

To get insight into the clonal expansion of T cell and B cell in brain lesion of LUAD patients, V(D)J rearrangements within the BCR/ TCRβ locus are PCR-amplified with V- and J-gene specific primers and the sequence of the CDR3 was determined using high-throughput sequencing of genomic DNA isolated from 3 patients with single-cell RNAseq data available. The abundance of individual T/B cell clonotypes is determined by combining templates from molecules with the same TCRB/BCR V-gene family (Figure 6A,B; supplemental fig.6A,B), J-gene segment(Figure 6A; supplemental fig.6A), and in-frameCDR3 amino acid sequence (Figure 6D; supplemental fig.6A,C). Among the brain lesions from three patients, atotal of 3,632,708 templates across 47,702 unique clonotypes were detected, with individual abundances ranging from 8,919 to 118,546 templates per clonotype(supplemental table3).

**Figure 6.**
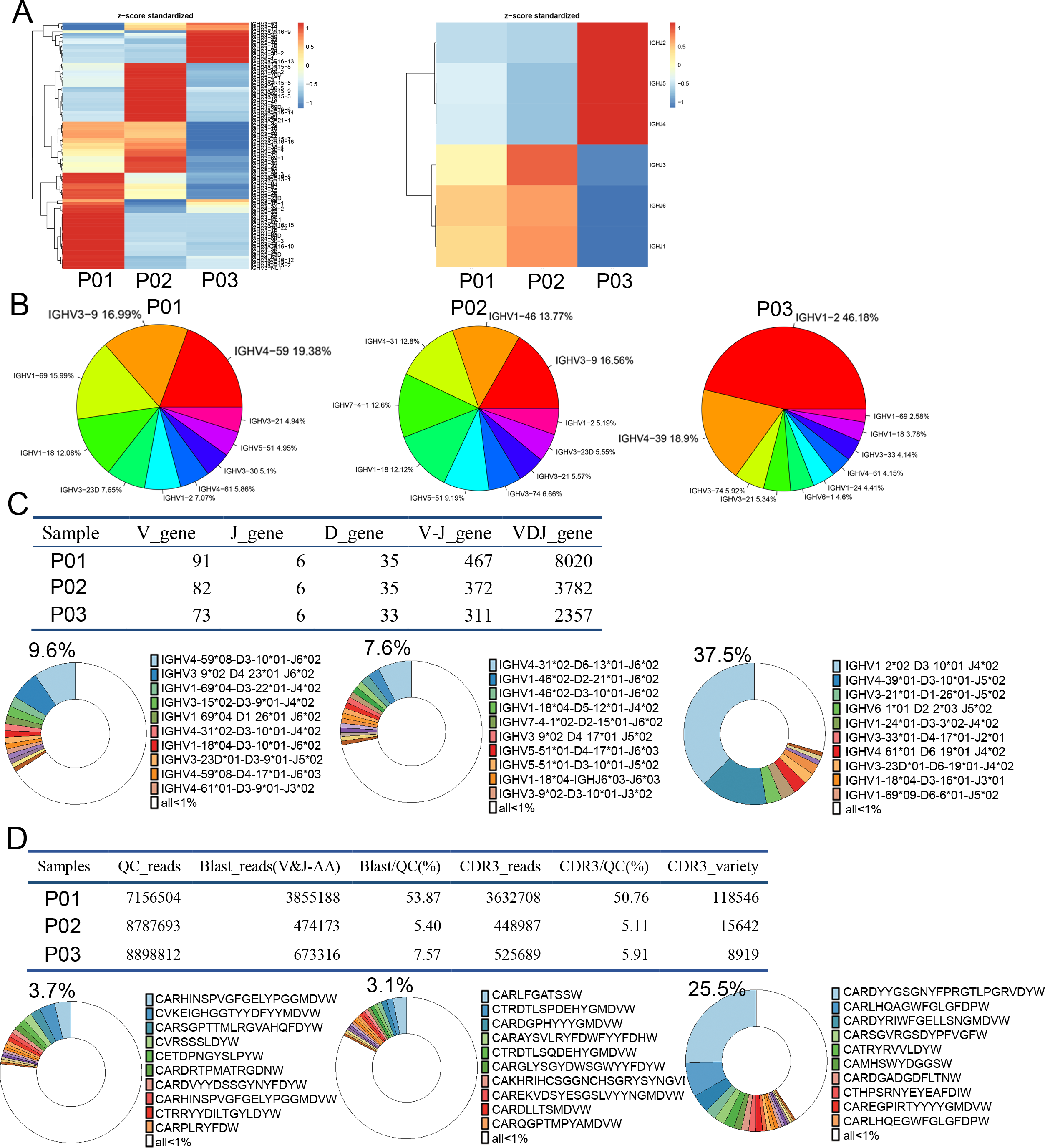
B cell clonality analysis of brain metastasis of LUADreveals diverse CDR3 sequence variety and clonal expansion. (A). Heatmap showing the clustering based on the frequency of V gene (left panel) and J gene (right panel). (B) Frequency of V gene usage for BCR clones across threedifferent patients. (C) Analysis of BCR clone diversity based on BCR VDJ rearrangement. The number of VDJ gene combinations are counted in threedifferent patients (top panel). The pie graph shows the frequency of representative VDJ combinations that account for more than 1% of the total.(D)Analysis of BCR clone diversity based on specific CDR3 sequence variety. The number of specific CDR3 sequences were counted in threedifferent patients (top panel). The pie graph shows the frequency of specific CDR3 sequences that account for more than 1% of the total.

In both CD3+ TCR and CD20+ BCR repertoires, the clonality (average 0.57 for BCR and 0.43 for TCR) in brain lesion of LUAD patients was generally higher than that observed in the primary tumor of a cohort of 35 NSCLC patients(average 0.07 for BCR and 0.3 for TCR)and also that of 35 standard healthy donor blood specimens (average 0.06 for BCR and 0.29 for TCR)(supplemental table 3)^51^. The heterogeneity of BCR genes across three patients also showed by the clustering of V or J gene usage (Figure 6A, B),diverse frequency of V genes (Figure 6B) or VDJ rearrangement (Figure 6C).

Three hundred BCR and 321 TCRβ clonotypes are present with an abundance higher than 0.1% in brain lesions (supplemental table 3). Strikingly, in one patient, one clonotype with BCR CDR3 sequence “CARDYYGSGNYFPRGTLPGRVDYW” accounts for 25.5% of the total clonotypes (Figure 6D). On the other hand, the TCRβ with CDR3 sequence” CASSFVTGGRTEAF” accounts for 17.2% of total TCRclonotypes in one patient (supplemental fig.6D, middle panel). This particular CDR3 sequence and also a lot of other high abundant CDR3 sequences share common motifs with a variety of virus reacting TCRs, including Cytomegalovirus (CMV), HIV-1, influenza (data not shown). The expansion of some of T cell clones are possibly not targeting the tumor-specific antigen but instead virus^52^, since the CD39 expression is also absent in majority of the T cells found in brain lesions (supplemental fig.2).

#### Conclusion

Despite the low number of T and B cells in brain metastases of LUAD patients, clonal expansion of T and B cells are apparent across the same three patients we already performed single cell RNAseq.

## Discussion

Brain metastases represent an unmet medical need in current oncologic care. Given the poor prognosis of patients with brain metastases, particularly in lung adenocarcinoma patients, there is critical need to improve our understanding of the mechanisms underlying the pathogenesis as well as to identify novel targets for immune therapies ^53–55^.

Advances in singlecell RNA sequencing (scRNA-seq) have allowed for a comprehensive analysis of intra-tumoral heterogeneity and tumor immune microenvironment in various cancer types at primary sites ^38,39,43,49,56–58^. But not much have been done on metastatic lesions, particularly brain metastases of different cancer types. Brain has been viewed as an immune-privileged organ, but accumulating evidences start to reveal the unique feature of brain immune microenvironment in obesity, neural degenerative disease and multiple sclerosis^59,60^. In order to understand the tumor immune microenvironment in brain lesion, we performed scRNA-seq on surgically resected brain lesionfrom three treatment-naïve LUAD patients and one triple negative breast cancer patient. We identified and profiling the different stromal cell lineages that accumulated in metastatic lesions, including immune cells (myeloid, T, and B cells), oligodendrocytes, endothelial cells, and pericytes. The expansion of macrophage and limited T and B cell infiltration is a common feature of these brain lesions. By comparing our scRNAseq data with published datasets derived from early and late-stage primary LUAD tumors^38,39^, we found that this compromised T cell response is unique to brain lesions, while the related abundant of T lymphocyte infiltration was found in primary LUAD patients. We also found the tumor immune microenvironment of brain metastases is shaped both by tumor intrinsic property that being inherent of their organ of origin (Lung and breast) and the stromal environment of their metastatic site (brain). First, when comparing with the tumor arising from brain, e.g., glioblastoma, molecular signature of stromal cell derive from lung and breast brain metastases are very distinct from their counterpart in glioblastoma (Figure 5B,C). This data indicates the tumor of origin play a significant role in determining their tumor microenvironment. On the other hand, when comparing with primary lung cancer (both early and later stage), the molecular signature of stromal cell in brain lesion of LUAD patients are well separated from their counterpart in primary lung tumor. This data indicates the strong influence of metastatic sites in shaping the tumor microenvironment. Based on these findings, we defined the unique molecular signature of stromal cells within brain lesions are likely attributed to the origin of tumor cells as well as its metastatic location.

Pro-inflammatory macrophage expansion and T cell dysfunction are two known microenvironment factor that contribute to immune escape^41,61,62^. Macrophages formerly act as scavengers to maintain tissue homeostasis. Mounting data suggest macrophages are abundant in the tumor microenvironment of brain tumors, when educated by cancer cells, they polarized to function as immune suppress cells^63–67^. We performed unbiased single-cell transcriptomic analysis of myeloid cells to demonstrate that TAM in the brain metastasis has a unique alternative activation M2 signature and a lack of conventional T cell co-stimulatory factors. In addition, brain metastases associated macrophages also express unique sets of immunosuppressiveligands(CD276 and VTCN1) comparing with their primary tumor counterpart (PDL1, CD80, LILRB2/5). Despite the low number of T and B cells in brain metastases of both lung and breast cancer, abundant TCR and BCR repertoire were identified in the same three patients we already performed single cell RNAseq. The clonal expansion of T and B cells are apparent across these patients. Although Chongsathidkiet et al. proposed sequestration of T cells in bone marrow as a tumor-adaptive mode of lack of T cell in brain tumor^68^, it is still unclear to what extent this could apply for the brain metastases of lung and breast tumor^37,69^. Since we found the TAM in brain lesion express high level of T cell chemoattractant such as CCL3 and CCL4 (supplemental fig.2B)^70^, so it is unlikely the lack of T cells is due to the limited T cell trafficking. The general immune suppressive environment of the brain may contribute partly to the lack of T/B cells in the three type of brain tumor we examined^63^. By profiling the known immune co-stimulatory factors in TAM in brain metastases, we find lack of expression of conventional co-stimulatory factors such as PVR, TNFSF9, ICOSLG, CD40LG^71^.

In summary, we constructed the first tumor microenvironment landscape of brain metastases of LUAD patients and triple negative breast cancer by massive parallel single cell RNA sequencing. We revealed the unique transcriptomic feature of tumor stromal cells that distinct them from their counterpart in primary tumor. Expansion of immune suppressive macrophage and lack of T cell activation and expansion may explain the immune escape of the brain metastases. Our study also identified some potential targets for checkpoint inhibitors that could be further developed as novel immunotherapy that target brain metastases. Further investigation about detail mechanisms how disseminated tumor cells interact with stromal cells in brain to shape their tumor microenvironment will significantly strengthen our understanding the pathogenesis of brain metastases. Comprehensive understanding of the composition and evolution of immune microenvironment of brain metastases will guide the design of new therapeutic strategies to target these deadly diseases^30^.

## Supporting information

Supplemental Table1 Patient information for priamry and brain metastases

Supplemental table 2. Known T cell regulatory ligand and their receptor

Supplemental table 3. Tcell-TCR and B cell-BCR VDJ rearrange ment and CDR3 variety

## Acknowledgements

This study was supported by National Key Research and Development Program of China (No.2017FYA0205302)to Wei-Lin Jin. National Natural Science Foundation of China (No. 81772661 and 81402081to Liang Wangand No. 81602193 to Wei Sun). J.D. is supported by a Postdoctoral Breakthrough award by the DoD’s BCRP (W81XWH-18-1-0028).

## Method

### Human Specimens

Tumor tissues were obtained from patients undergoing brain resection surgery at the Tangdu Hospital of Fourth Military Medical University (Xi’an, China) after obtaining informed consent. All protocols were reviewed and approved by the Institutional Review Board (IRB) at the Tangdu Hospital (TDLL-201709-20).

### Patient Clinical Characteristics

Samples were collected from 19 patients who were diagnosed with lung cancer and brain lesions were confirmed as lung adenocarcinoma (LUAD) subtype and 1 triple negative breast cancer (TNBC) patients with brain metastasis(Supplemental table 1). The 19 LUAD patients consisted of 11 males and 8 females and ranged from age 28 to 72 years old, with a median of 52 years old. The TNBC patient 44 was years old.

### Sample Collection and digestion for single cell RNAseq

From the 3 of the LUAD patients and 1 TNBC patient described above, brain tissue containing the metastatic lesions was collected under surgeries at Tangdu Hospital with informed consent under a protocol approved by Fourth Military Medical University Institutional Review Board. Tissue samples were placed in DMEM/RPMI media (Corning) on ice. Small chunk of the tissue was processed for immunohistochemistry analysis by quick frozen section. H&E slides were reviewed by a pathologist and confirmed as LUAD brain metastasis or TNBC brain metastasis. The remaining freshly dissected brain tissue was cut up and then enzymatically dissociated to obtain a suspension of cells using a Tumor Dissociation Kit (MiltenyiBiotec#130-095-929) by incubating at 37 ℃ for 15 minutes. Dissociated single cells were pelleted by centrifuged at 130g for 10 minutes after removing unlysed tissue fragments with a 20μm cell strainer. The cell pellet was then resuspended in Dulbecco’s phosphate-buffered saline (DPBS) containing 0.02% BSA. Cell health was assessed by trypan blue exclusion. Ensure that no less than 90% of the cell samples are alive for subsequent single-cell RNAseq (10X Genomics).

### Single cell capture, library preparation, and sequencing

Single-cell libraries were prepared using the Chromium 3′ v2 platform (10X Genomics) following the manufacturer’s protocol[1] by Capital Bio Technology Corporation and Novogene. Briefly, single cells were partitioned into Gel beads in Emulsion in the 10X Chromium Controller instrument followed by cell lysis and barcoded reverse transcription of RNA, amplification, shearing and 5′ adapter and sample index attachment. On average, ~10,000 cells were loaded on each channel that resulted in the recovery of ~7822 cells. Libraries were sequenced on Illumina HiSeq2500 instruments using paired-end sequencing (PE1 54 bp and PE2 66 bp). Each replicate was sequenced on one half of a HiSeq lane, at an initial depth of approximately 100 million reads.

### Alignment, barcode assignment and UMI counting

The Cell Ranger Single-Cell Software Suite was used to perform sample demultiplexing, barcode processing and single-cell 3′ gene counting. First, sample demultiplexing was performed based on the 8 bp sample index read to generate FASTQs for the Read1 and Read2 paired-end reads, as well as the 14 bp GemCode barcode. Then, Read1, which contains the cDNA insert, was aligned to an appropriate reference genome using STAR. Next, GemCode barcodes and UMIs were filtered. All of the known listed of barcodes that are 1-Hamming-distance away from an observed barcode are considered.

Cell barcodes were determined based on distribution of UMI counts. Number of reads that provide meaningful information is calculated as the product of four metrics: (1) valid barcodes; (2) valid UMI; (3) associated with a cell barcode; and (4) confidently mapped to exons. Samples processed from multiple channels can be combined by concatenating gene-cell-barcode matrices. In addition, sequencing data can be subsampled to obtain a given number of UMI counts per cell. For 3 LUAD patients, on average, 7,822 cells were sequenced for each patient, with 135,464 reads per cell and 2,532 genes per cells. For the one triple negative breast cancer patient, 6000 cells were sequenced, with 124,080 reads per cell and 575 genes per cells. Detailed parameter of sequencing quality are shown in Supplemental Figure 1.

### Single cell RNAseq data preprocessing

Gene expression datasets were processed with the R package of *Seurat* version 2. Cells were retained for downstream analysis based on the following quality measures: number of genes detected greater than 500, percentage of mitochondrial transcripts less than 5%. Raw UMI counts were normalized by dividing the number of UMIs of each gene by all UMIs of a given cell and then multiplied by 10,000. After normalization, confounding factors including the number of detected genes and proportions of mitochondrial transcripts were also regressed out. Highly variable genes used for canonical correlation analysis (CCA)with multiple groups must be highly variable in at least two datasets and having dispersion higher than 0.5 and normalized expression level between 0.0125 and 3. The top 10 PCs established with principal component analysis (PCA) were used to do clustering using Seurat. Cluster identities were assigned based on cluster gene markers as determined by *FindAllMarkers* function in Seuratand gene expression of known marker genes. Cluster markers were determined using *Wilcoxon* test as genes showing a minimum log fold expression change of 0.25 in at least a fraction of 25% of cells in the clusters.

Cells were typed by examining expression of known marker genes. In this analysis, cells were scored as detecting a marker gene if the cell contained a non-zero molecule count for that gene. Each cell was corrected for its detection rate (the fraction of total genes detected in that cell) and the marker detection rate was then averaged across cells of a cluster. Markers used to type cells included NCAM1, NCR1, NKG2 (NK-cells), GNLY, GZMA, GZMB, GMZM (cytotoxic T, NK), FOXP3(T-regulatory Cell), CTLA4, TIGIT, TNFRSF4, LAG3, PDCD1 (Exhausted T-cell), CD8, CD3, CD4 (T-cells), IL7R (Naive T-cells), CD19 (B-cells), CD79A, IGHG1, IGHA1, IGHM, JCHAIN, MZB1 (Plasma),ENPP3, KIT (Mast cells), IL3RA, LILRA4 (plasmacytoid DC), HLA-DR, FCGR3A, SPP1, AIF1, APOE, CD68, CD63, TREM2, CD14, ITGAM, (Monocytic Lineage),PECAM1, CLDN5, VWF, ESAM (Endothelia), ACTA2,PDGFRB, TAGLN, DCN (Pericyte), MAG, NKX6.2, OLIG1, OLIG2, PLP1, ERMN, MOG(Oligodendrocyte).

### Molecular signatures of microglia/macrophage

Microglia/macrophage belonging to clusters 3,5,9,10,16,19,20 (Figure 2B, D) were extracted from the 5,187 cells dataset and new *Seurat* objects were built using function setup for microglia, macrophage, monocyte and dendritic cells. Multiple datasets were integrated with *RunMultiCCA* function in *Seurat* package. The marker genes were identified by using the *Wilcoxon*test [p-value< 0.001, log2(fold change)>1], which were also used for constructing a gene expression matrix for heatmap. R tSNE with default parameters in *Seurat* package was used to build a 2D map for group visualization. Each dot represented a single cell and each cluster was colored with scheme consistent with other figures. Heatmaps in Figure 4 was generated by *ComplexHeatmap* R package.

### Process data from published scRNA

Early LUAD from GSE97168, [2]; Late lung cancer [3] and Glioblastoma datasets from GSE103224[4].

### RNA isolation and bulk RNAseq

Total RNA was extracted using Qiagen Total RNA Isolation Kit, and the integrity and concentration were evaluated using an Agilent 2100 Bioanalyzer (Agilent Technologies, USA). Total RNA was purified to obtain mRNA using oligo (dT) magnetic beads. The cDNA libraries were constructed using Illumina TruSeq RNA Sample Prep Kit (Illumina, USA) according to the manufacturer’s protocol. The quality of the constructed cDNA libraries was examined on an Agilent Technologies 2100 382 Bioanalyzer prior to sequencing with Illumina HiSeq™ 2500 at Shanghai OE Biotechnology Co., Ltd. (http://www.oebiotech.com/). Bcl to fastq conversion was done with Illumina bcl2fastq software and demultiplexed into individual files of each sample with self-written scripts in Perl according to the indices for pooling. Low-quality reads were filtered out with Trimmomatic software with default parameters and then clean reads were aligned to the indexed human GRCh38 genome reference with HISAT2 software to generate the SAM files, which were finally converted into sorted BAM files with SAMtools software. Alignment of reads into target genes was performed by using the Stringtie software [5], and gene counts were calculated using the Stringtie/prepDE.py with default options. Log10-transformed TPM (transcript per million reads) was used to calculate and compare gene expression differences between different samples. In this study, p < 0.05 and a fold change (FC) > 2 (|log2FC|>1) in at least one treatment were set as thresholds to define the significance of gene expression differences between two treatments (Supplementary fig.5A). The Blast2GO software was applied to get the annotation results of unigenes in the Gene Ontology (GO) database. The pathways were also annotated according to the KEGG database.

#### TCGA Analysis

Tumor RNAseq counts data from 11 patients of stage I lung adenocarcinoma (early LUAD) and 11 patients with stage IV (late LUAD) patients were downloaded from Genomic Data Commons Portal https://gdc-portal.nci.nih.gov/ (supplemental table 1). Normalized log10-transformed expression (TPM) was used for downstream analysis.

### TCR or BCR repertoire sequencing

Immunosequencing of the CDR3 regions of human TCRβ chains and BCR was performed by Oebiotech (Oebiotech,Shanghai, China). Briefly, two-round PCR was performed for TCR and BCR library preparation as described[6], followed by high-throughput sequencing. Sequences were collapsed and filtered in order to identify and quantitate the absolute abundance of each unique TCRβ CDR3 region or BCR CDR3 region for further analysis as previously described[7, 8]

### Statistical Analyses of TCR-β Sequencing

Clonality was defined as 1-the normalized Shannon entropy[9, 10].Clonality values range from 0 to 1 and describe the shape of the frequency distribution: clonality values approaching 0 indicate a very even distribution of frequencies, whereas values approaching 1 indicate an increasingly asymmetric distribution in which a few clones are present at high frequencies. Clonality between experimental groups was compared using a two-tailed Wilcoxon Rank Sum test.

### Immunohistochemical analysis

Lung adenocarcinoma with lung tissue microarray, containing 48 cases adenocarcinoma and matched cancer adjacent or adjacent normal lung tissue was purchased from Alenobio (Cat#LC10013b) and brain lesions from 15 patients with LUAD were surgical resected and formalin fixed and paraffin embedded. Immunohistochemistry using standard immunoperoxidase staining was performed on formalin-fixed paraffin-embedded tissue sections (5 μm thick) from specimens of each of the tumor resections. Briefly, we used 3 × 3 min cycles of de-paraffinization in xylene, 2 × 1 min cycles of dehydration in 100% ethanol, 2 × 1 min cycles of dehydration in 95% ethanol, and a 1-min cycle of dehydration in 70% ethanol. Slides were then washed in water. We used 0.01 M citrate buffer (pH 6) for antigen retrieval in a microwaved pressure cooker for 20 min. We then washed the slides three times in phosphage-buffered saline (PBS) after cooling for 30 min. We quenched endogenous peroxidase in 3% hydrogen peroxide in PBS for 10 min, washed three times in PBS, and blocked with 10% goat serum for 25 min. We then incubated the slides with primary antibodies for 90 min at room temperature. We used the following primary antibodies: mouse anti-CD20(ORIGENE, ZM-0039, 1:100 dilution), anti-CD3(ORIGENE, ZM-0417,1:100 dilution), anti-CD68(ORIGENE, ZM-0464, 1:100 dilution). After washing three times in PBS, we incubated the slides with biotinylated goat anti-rabbit secondary antibody (Vector Laboratories, 1:200 dilution) for 30 min at room temperature, followed by additional PBS washing, 30-min incubation with ABC peroxidase reagent, development in DAB-peroxidase substrate solution (DAKO), and counter-staining in hematoxylin. Images were taken at 10×5 by LEICA DMC 4500 and analyzed using Image-Pro Plus 6.0.

## Supplemental table description

Table S1. Detail clinical information of 19 Lung adenocarcinoma (LUAD) patients with brain metastases and 1 triple negative breast cancer patient with brain metastases. Also the 32 primary LUAD patients from lung tissue microarray purchased from Alenobio (Cat#LC10013b) and primary LUAD patients from TCGA database.

Table S2. List of known T cell regulatory ligands and their receptors used to make Figure 4D, E.

Table S3. Detailed sequences information for TCR and BCR VDJ rearrangement and CDR3 variety of three patients and T/B cell clonality index in 19 primary LUAD patient [11] and brain metastases of 3 LUAD patients.

---

**Supplemental fig.1.**
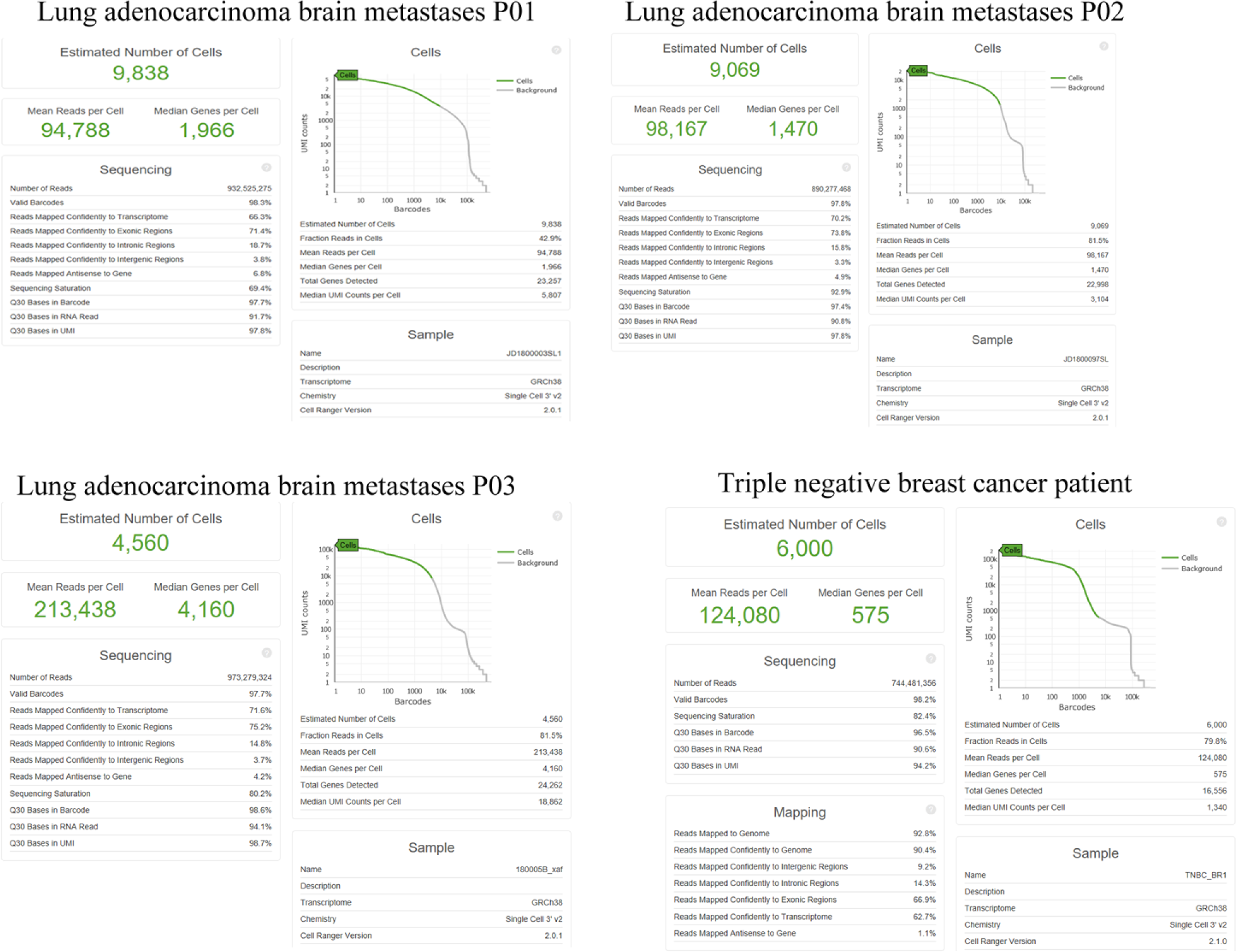
Web summary from the raw sequencing data processing after 10X genomic Cell Ranger Algorithms. Number of cells detected, the mean reads per cell, and the median genes detected per cell are prominently displayed near the top of each panel. Data from three lung adenocarcinoma (LUAD) patients (P01, P02, P03) and one triple negative breast cancer patient (TNBC).

**Supplemental fig.2.**
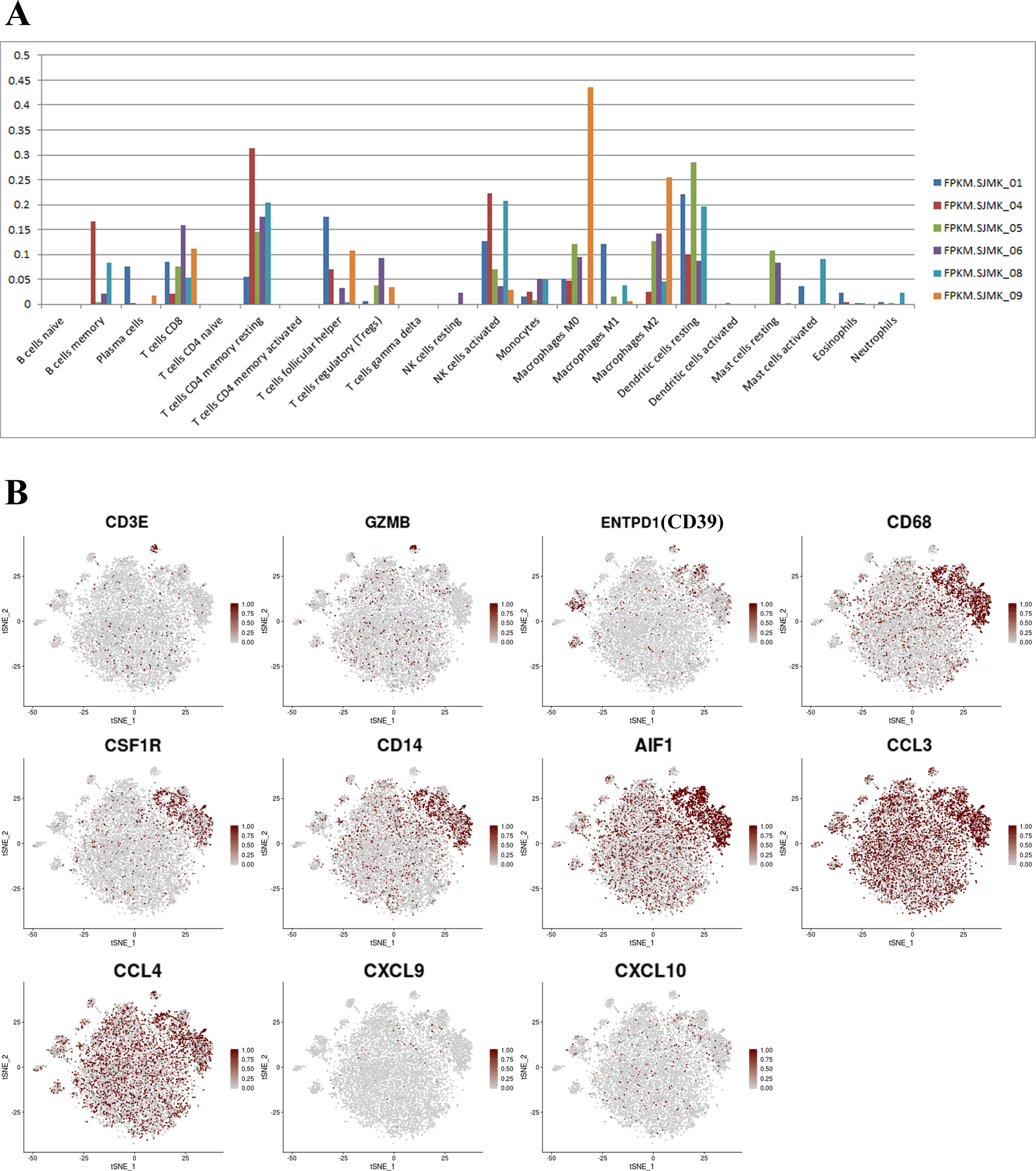
(A) Relative leukocyte fractions evaluated by CIBERSORT in our bulk RNAseq data to infer relative RNA fractions from 22 leukocyte subsets (LM22 signature) in each patient represented with different colors. (B) tSNE plot color-coded for expression (low to high: gray to dark red) of marker genes. Clusters are indicated in figure 1B. Cytotoxic T cells:CD3E, GMZB and ENTPDA(CD39); Macrophages: CD68, CSFR1, CD14, AIF1, CCL3, CCL4, CXCL9 and CXCL10.

**Supplemental fig.3.**
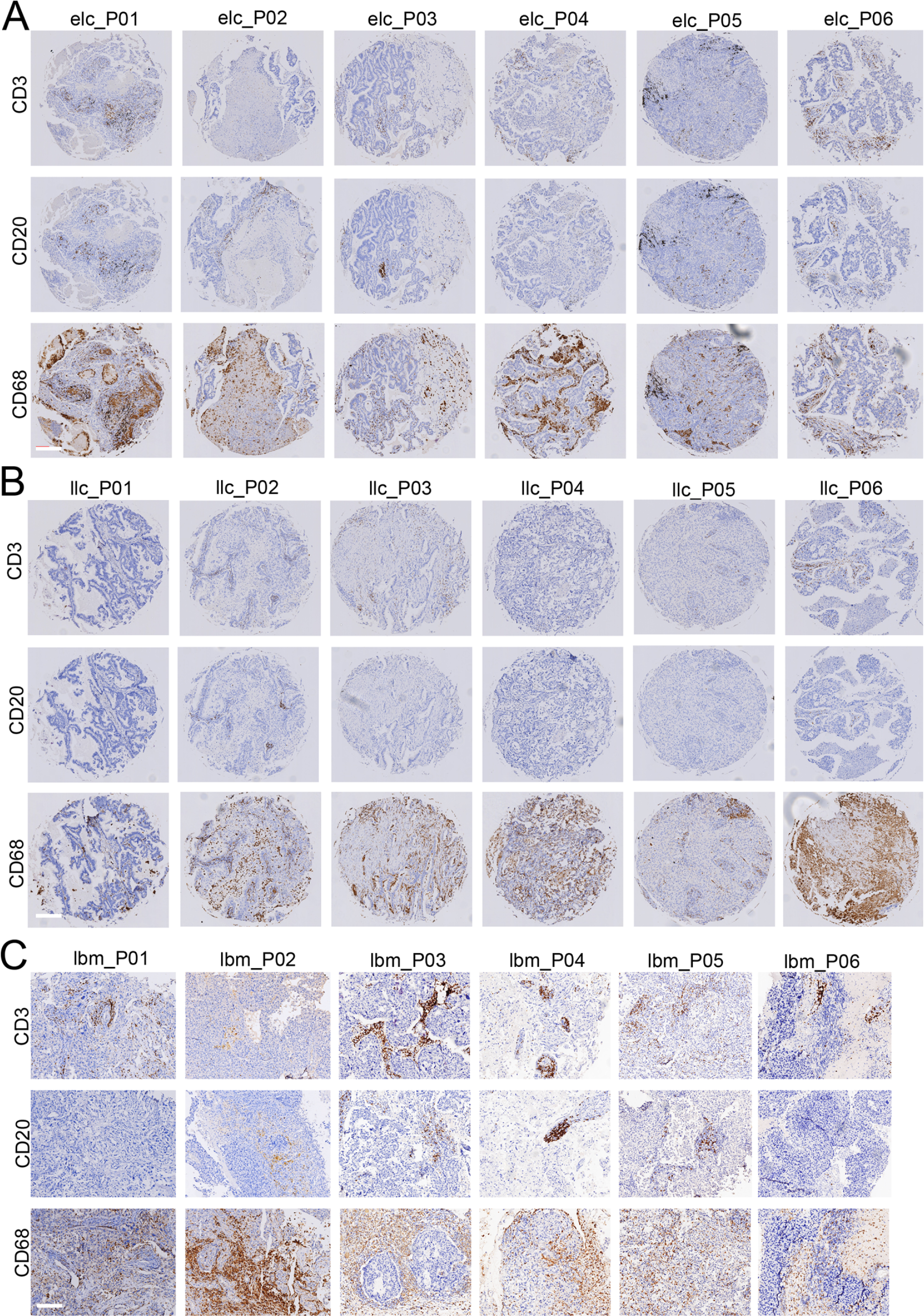
Immunohistochemistry assessment of formalin-fixed paraffin embedded archival tissue samples (×200 magnification). For each patient, samples were stained with CD3 (T cell marker), CD20 (B cell marker) and CD68 (macrophage marker), respectively. Representative images shown as (A) six early stage primary LUADs (elc, stage IA/B); (B) six late stage primary LUADs (llc, stage IV) and six brain metastatic lesions from treatment naïve LUAD patients (lbm). Scale bar:200um.

**Supplemental fig.4.**
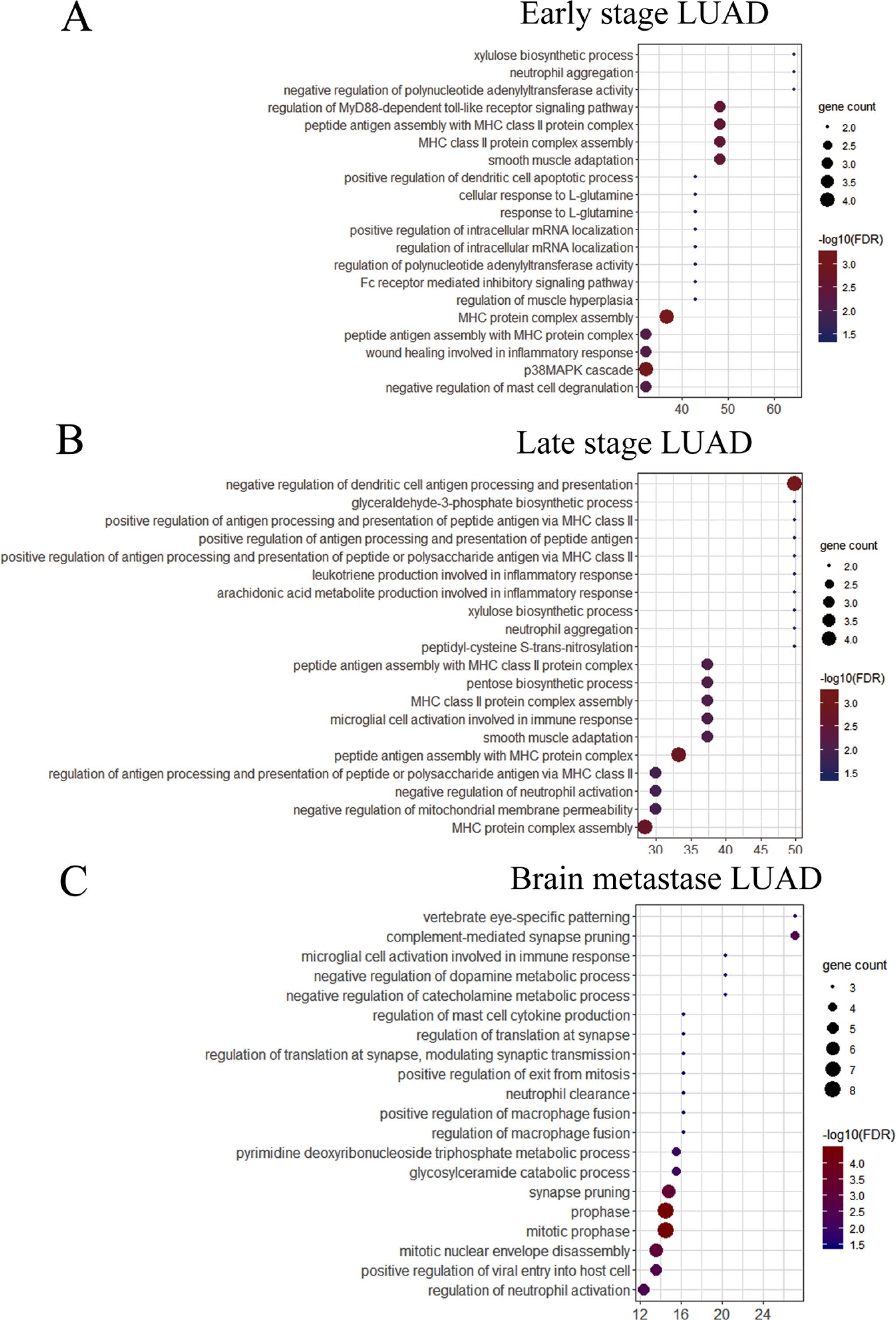
Gene ontology (biological process) analysis of the differentially expressed genes enriched in TAM clusters in different stages of LUAD:(A) Early stage primary LUAD (elc, cluster 9); (B) Late stage primary LUAD (llc, cluster 5); (C) LUAD brain metastases (lbm, cluster 20). Top20 over-represented GO terms with higher enrichment scores and FDR (false discovery rate) significance less than 0.05are shown.

**Supplemental fig.5.**
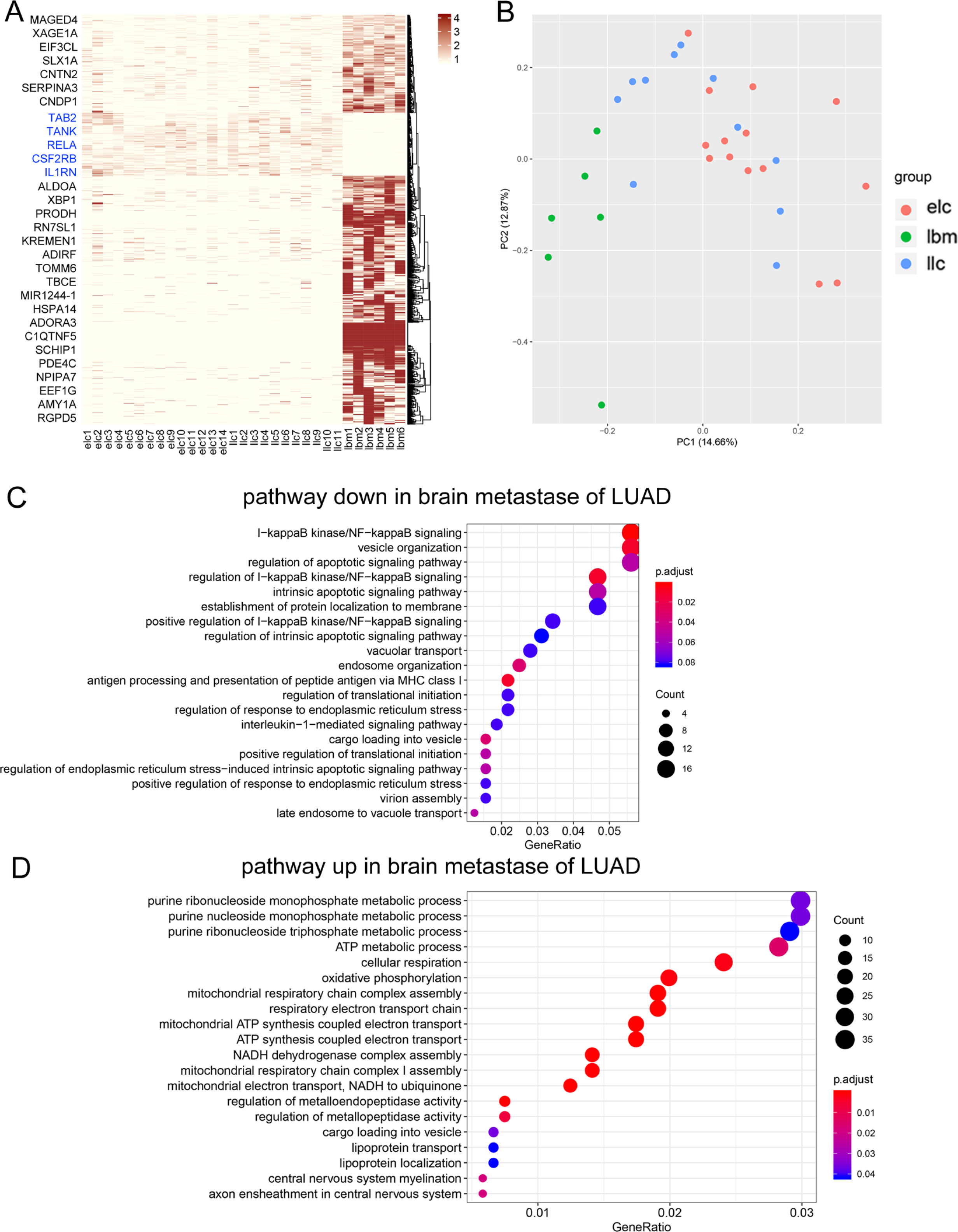
(A) Unsupervised hierarchical clustering heatmap of gene expressions (red, high relative expression; blue, low relative expression) of three patient cohorts: early stage IA/B primary LUAD patients (elc, n=14), late stage V primary LUAD patients (llc, n=11) and LUAD patients with brain metastases (lbm, n=6). Differentially expressed genes (n=1,999 higher and 362 lower in lbm) with fold change>±2, and p<0.05 (Wilcoxon sum rank test) were used across the cohorts. (B) Principle component analysis (PCA) of gene expression profiles from different stages of LUAD patients. (C, D) Gene ontology (GO Biological process) enrichment analysis of the differentially expressed genes downregulated (C) or upregulated (D)in LUAD brain metastases (lbm) compared with both early stage primary LUAD (elc) and late stage primary LUAD (llc). Top20 over-represented pathways with higher gene ratio (Gene Ratio) and adjusted p-value less than 0.05 are shown.

**Supplemental fig.6.**
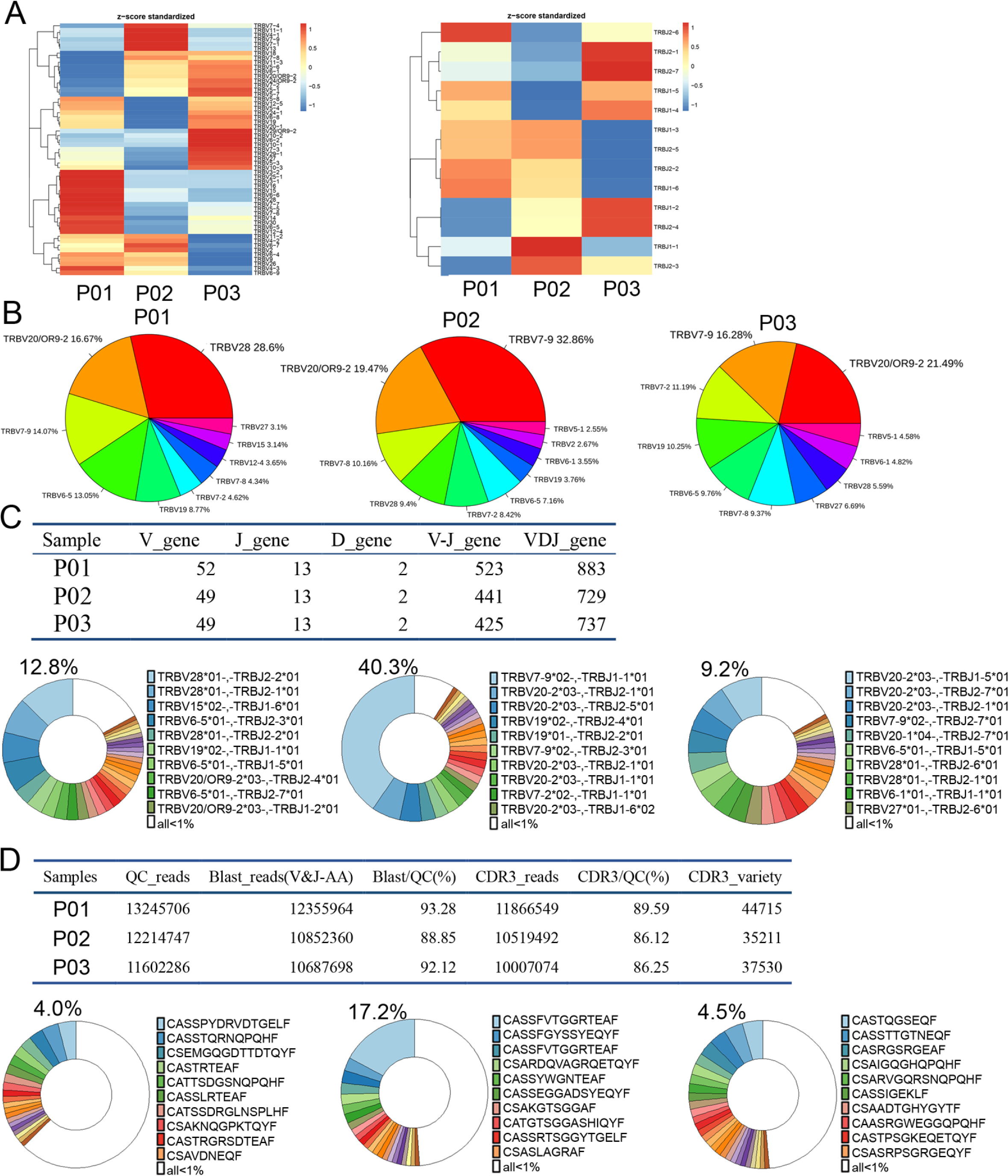
(A) Heatmap showing the clustering based on the frequency of V gene (left panel) and J gene (right panel). (B) Frequency of V gene usage for TCRβ clones across three different patients. (C) Analysis of TCR clone diversity based on TCR VDJ rearrangement. The number of VDJ gene combinations are counted in three different patients (top panel). The pie graph shows the frequency of representative VDJ combinations that account for more than 1% of total. (D) Analysis of TCR clone diversity based on specific CDR3 sequence variety. The number of specific CDR3 sequences were counted in three different patients (top panel). The pie graph shows the frequency of specific CDR3 sequences that account for more than 1% of total.

## Reference

1. Arvold, N.D., et al. Updates in the management of brain metastases. Neuro Oncol18, 1043–1065 (2016).

2. Johnson, J.D. & Young, B. Demographics of brain metastasis. Neurosurg Clin N Am7, 337–344 (1996).

3. Barnholtz-Sloan, J.S., et al. Incidence proportions of brain metastases in patients diagnosed (1973 to 2001) in the Metropolitan Detroit Cancer Surveillance System. J Clin Oncol22, 2865–2872 (2004).

4. Schouten, L.J., Rutten, J., Huveneers, H.A. & Twijnstra, A. Incidence of brain metastases in a cohort of patients with carcinoma of the breast, colon, kidney, and lung and melanoma. Cancer94, 2698–2705 (2002).

5. Achrol, A.S., et al. Brain metastases. Nat Rev Dis Primers5, 5 (2019).

6. Steeg, P.S., Zimmer, A. & Gril, B. Therapeutics for Brain Metastases, v3. Clin Cancer Res22, 5953–5955 (2016).

7. Chan, B.A. & Hughes, B.G. Targeted therapy for non-small cell lung cancer: current standards and the promise of the future. Transl Lung Cancer Res4, 36–54 (2015).

8. Zhang, C., Leighl, N.B., Wu, Y.L. & Zhong, W.Z. Emerging therapies for non-small cell lung cancer. J Hematol Oncol12, 45 (2019).

9. Shaw, A.T. & Engelman, J.A. Ceritinib in ALK-rearranged non-small-cell lung cancer. N Engl J Med370, 2537–2539 (2014).

10. Shanker, M., Willcutts, D., Roth, J.A. & Ramesh, R. Drug resistance in lung cancer. Lung Cancer (Auckl)1, 23–36 (2010).

11. Jamal-Hanjani, M., et al. Tracking the Evolution of Non-Small-Cell Lung Cancer. N Engl J Med376, 2109–2121 (2017).

12. Sequist, L.V., et al. Genotypic and histological evolution of lung cancers acquiring resistance to EGFR inhibitors. Sci Transl Med3, 75ra26 (2011).

13. Pakkala, S. & Ramalingam, S.S. Personalized therapy for lung cancer: striking a moving target. JCI Insight3(2018).

14. Crequit, P., et al. Comparative efficacy and safety of second-line treatments for advanced non-small cell lung cancer with wild-type or unknown status for epidermal growth factor receptor: a systematic review and network meta-analysis. BMC Med15, 193 (2017).

15. Hodi, F.S., et al. Improved survival with ipilimumab in patients with metastatic melanoma. N Engl J Med363, 711–723 (2010).

16. Robert, C., et al. Pembrolizumab versus Ipilimumab in Advanced Melanoma. N Engl J Med372, 2521–2532 (2015).

17. Reck, M., et al. Pembrolizumab versus Chemotherapy for PD-L1-Positive Non-Small-Cell Lung Cancer. N Engl J Med375, 1823–1833 (2016).

18. Borghaei, H., et al. Nivolumab versus Docetaxel in Advanced Nonsquamous Non-Small-Cell Lung Cancer. N Engl J Med373, 1627–1639 (2015).

19. Brahmer, J., et al. Nivolumab versus Docetaxel in Advanced Squamous-Cell Non-Small-Cell Lung Cancer. N Engl J Med373, 123–135 (2015).

20. Herbst, R.S., et al. Pembrolizumab versus docetaxel for previously treated, PD-L1-positive, advanced non-small-cell lung cancer (KEYNOTE-010): a randomised controlled trial. Lancet387, 1540–1550 (2016).

21. Schartz, N.E., et al. Complete regression of a previously untreated melanoma brain metastasis with ipilimumab. Melanoma Res20, 247–250 (2010).

22. Hodi, F.S., et al. CTLA-4 blockade with ipilimumab induces significant clinical benefit in a female with melanoma metastases to the CNS. Nat Clin Pract Oncol5, 557–561 (2008).

23. Queirolo, P., et al. Efficacy and safety of ipilimumab in patients with advanced melanoma and brain metastases. J Neurooncol118, 109–116 (2014).

24. Konstantinou, M.P., et al. Ipilimumab in melanoma patients with brain metastasis: a retro-spective multicentre evaluation of thirty-eight patients. Acta Derm Venereol94, 45–49 (2014).

25. Goldberg, S.B., et al. Pembrolizumab for patients with melanoma or non-small-cell lung cancer and untreated brain metastases: early analysis of a non-randomised, open-label, phase 2 trial. Lancet Oncol17, 976–983 (2016).

26. Dudnik, E., et al. Intracranial response to nivolumab in NSCLC patients with untreated or progressing CNS metastases. Lung Cancer98, 114–117 (2016).

27. Johanns, T., Waqar, S.N. & Morgensztern D. Immune checkpoint inhibition in patients with brain metastases. Ann Transl Med4, S9 (2016).

28. Doroshow, D.B., et al. Immunotherapy in Non-Small Cell Lung Cancer: Facts and Hopes. Clin Cancer Res25, 4592–4602 (2019).

29. Cho, J.H. Immunotherapy for Non-small-cell Lung Cancer: Current Status and Future Obstacles. Immune Netw17, 378–391 (2017).

30. Binnewies, M., et al. Understanding the tumor immune microenvironment (TIME) for effective therapy. Nat Med24, 541–550 (2018).

31. Amir el, A.D., et al. viSNE enables visualization of high dimensional single-cell data and reveals phenotypic heterogeneity of leukemia. Nat Biotechnol31, 545–552 (2013).

32. Newman, A.M., et al. Robust enumeration of cell subsets from tissue expression profiles. Nat Methods12, 453–457 (2015).

33. Finotello, F. & Trajanoski, Z. Quantifying tumor-infiltrating immune cells from transcriptomics data. Cancer Immunol Immunother67, 1031–1040 (2018).

34. Al-Shibli, K.I., et al. Prognostic effect of epithelial and stromal lymphocyte infiltration in non-small cell lung cancer. Clin Cancer Res14, 5220–5227 (2008).

35. Salmon, H., et al. Matrix architecture defines the preferential localization and migration of T cells into the stroma of human lung tumors. J Clin Invest122, 899–910 (2012).

36. Turley, S.J., Cremasco, V. & Astarita, J. L. Immunological hallmarks of stromal cells in the tumour microenvironment. Nat Rev Immunol15, 669–682 (2015).

37. Anderson, K.G., Stromnes, I.M. & Greenberg, P. D. Obstacles Posed by the Tumor Microenvironment to T cell Activity: A Case for Synergistic Therapies. Cancer Cell31, 311–325 (2017).

38. Lavin, Y., et al. Innate Immune Landscape in Early Lung Adenocarcinoma by Paired Single-Cell Analyses. Cell169, 750–765 e717 (2017).

39. Lambrechts, D., et al. Phenotype molding of stromal cells in the lung tumor microenvironment. Nat Med24, 1277–1289 (2018).

40. Engblom, C., Pfirschke, C. & Pittet, M. J. The role of myeloid cells in cancer therapies. Nat Rev Cancer16, 447–462 (2016).

41. Grivennikov, S.I., Greten, F.R. & Karin M. Immunity, inflammation, and cancer. Cell140, 883–899 (2010).

42. Peranzoni, E., et al. Macrophages impede CD8 T cells from reaching tumor cells and limit the efficacy of anti-PD-1 treatment. Proc Natl Acad Sci U S A115, E4041–E4050 (2018).

43. Azizi, E., et al. Single-Cell Map of Diverse Immune Phenotypes in the Breast Tumor Microenvironment. Cell174, 1293–1308 e1236 (2018).

44. Murray, P. J. Macrophage Polarization. Annu Rev Physiol79, 541–566 (2017).

45. Muller, S., et al. Single-cell profiling of human gliomas reveals macrophage ontogeny as a basis for regional differences in macrophage activation in the tumor microenvironment. Genome Biol18, 234 (2017).

46. Xue, J., et al. Transcriptome-based network analysis reveals a spectrum model of human macrophage activation. Immunity40, 274–288 (2014).

47. Gosselin, D., et al. An environment-dependent transcriptional network specifies human microglia identity. Science356(2017).

48. Guo, X., et al. Global characterization of T cells in non-small-cell lung cancer by single-cell sequencing. Nat Med24, 978–985 (2018).

49. Yuan, J., et al. Single-cell transcriptome analysis of lineage diversity in high-grade glioma. Genome Med10, 57 (2018).

50. Wellenstein, M.D. & de Visser, K. E. Cancer-Cell-Intrinsic Mechanisms Shaping the Tumor Immune Landscape. Immunity48, 399–416 (2018).

51. Zhu, W., et al. A high density of tertiary lymphoid structure B cells in lung tumors is associated with increased CD4(+) T cell receptor repertoire clonality. Oncoimmunology4, e1051922 (2015).

52. Simoni, Y., et al. Bystander CD8(+) T cells are abundant and phenotypically distinct in human tumour infiltrates. Nature557, 575–579 (2018).

53. Dagogo-Jack, I., Carter, S.L. & Brastianos, P. K. Brain Metastasis: Clinical Implications of Branched Evolution. Trends Cancer2, 332–337 (2016).

54. Franchino, F., Ruda, R. & Soffietti R. Mechanisms and Therapy for Cancer Metastasis to the Brain. Front Oncol8, 161 (2018).

55. Di Giacomo, A.M., et al. Immunotherapy of brain metastases: breaking a “dogma”. J Exp Clin Cancer Res38, 419 (2019).

56. Baslan, T. & Hicks J. Unravelling biology and shifting paradigms in cancer with single-cell sequencing. Nat Rev Cancer17, 557–569 (2017).

57. Lawson, D.A., Kessenbrock, K., Davis, R.T., Pervolarakis, N. & Werb Z. Tumour heterogeneity and metastasis at single-cell resolution. Nat Cell Biol20, 1349–1360 (2018).

58. Tirosh, I., et al. Dissecting the multicellular ecosystem of metastatic melanoma by single-cell RNA-seq. Science352, 189–196 (2016).

59. Carson, M.J., Doose, J.M., Melchior, B., Schmid, C.D. & Ploix, C.C. CNS immune privilege: hiding in plain sight. Immunol Rev213, 48–65 (2006).

60. Negi, N. & Das, B.K. CNS: Not an immunoprivilaged site anymore but a virtual secondary lymphoid organ. Int Rev Immunol37, 57–68 (2018).

61. Dunn, G.P., Old, L.J. & Schreiber, R. D. The three Es of cancer immunoediting. Annu Rev Immunol22, 329–360 (2004).

62. Thommen, D.S. & Schumacher, T.N. T Cell Dysfunction in Cancer. Cancer Cell33, 547–562 (2018).

63. Quail, D.F. & Joyce, J. A. The Microenvironmental Landscape of Brain Tumors. Cancer Cell31, 326–341 (2017).

64. Sica, A. & Bronte, V. Altered macrophage differentiation and immune dysfunction in tumor development. J Clin Invest117, 1155–1166 (2007).

65. Pathria, P., Louis, T.L. & Varner, J. A. Targeting Tumor-Associated Macrophages in Cancer. Trends Immunol40, 310–327 (2019).

66. Filipazzi, P., Huber, V. & Rivoltini, L. Phenotype, function and clinical implications of myeloid-derived suppressor cells in cancer patients. Cancer Immunol Immunother61, 255–263 (2012).

67. Sica, A., et al. Macrophage polarization in tumour progression. Semin Cancer Biol18, 349–355 (2008).

68. Chongsathidkiet, P., et al. Sequestration of T cells in bone marrow in the setting of glioblastoma and other intracranial tumors. Nat Med24, 1459–1468 (2018).

69. Valiente, M., et al. The Evolving Landscape of Brain Metastasis. Trends Cancer4, 176–196 (2018).

70. Castellino, F., et al. Chemokines enhance immunity by guiding naive CD8+ T cells to sites of CD4+ T cell-dendritic cell interaction. Nature440, 890–895 (2006).

71. Chen, L. & Flies, D. B. Molecular mechanisms of T cell co-stimulation and co-inhibition. Nat Rev Immunol13, 227–242 (2013).

## Reference

1. Zheng, G.X., et al., Massively parallel digital transcriptional profiling of single cells. Nat Commun, 2017.8: p. 14049.

2. Lavin, Y., et al., Innate Immune Landscape in Early Lung Adenocarcinoma by Paired Single-Cell Analyses. Cell, 2017.169(4): p. 750–765 e17.

3. Lambrechts, D., et al., Phenotype molding of stromal cells in the lung tumor microenvironment. Nat Med, 2018.24(8): p. 1277–1289.

4. Yuan, J., et al., Single-cell transcriptome analysis of lineage diversity in high-grade glioma. Genome Med, 2018.10(1): p. 57.

5. Pertea, M., et al., Transcript-level expression analysis of RNA-seq experiments with HISAT, StringTie and Ballgown. Nat Protoc, 2016.11(9): p. 1650–67.

6. Chen, Y., et al., High-throughput T cell receptor sequencing reveals distinct repertoires between tumor and adjacent non-tumor tissues in HBV-associated HCC. Oncoimmunology, 2016.5(10): p. e1219010.

7. Robins, H.S., et al., Comprehensive assessment of T-cell receptor beta-chain diversity in alphabeta T cells. Blood, 2009.114(19): p. 4099–107.

8. Carlson, C.S., et al., Using synthetic templates to design an unbiased multiplex PCR assay. Nat Commun, 2013.4: p. 2680.

9. Kirsch, I., M. Vignali, and H. Robins, T-cell receptor profiling in cancer. Mol Oncol, 2015.9(10): p. 2063–70.

10. Jia, Q., et al., Diversity index of mucosal resident T lymphocyte repertoire predicts clinical prognosis in gastric cancer. Oncoimmunology, 2015.4(4): p. e1001230.

11. Zhu, W., et al., A high density of tertiary lymphoid structure B cells in lung tumors is associated with increased CD4(+) T cell receptor repertoire clonality. Oncoimmunology, 2015.4(12): p. e1051922.

